# Intragenomic variation in mutation biases causes underestimation of selection on synonymous codon usage

**DOI:** 10.1101/2021.10.29.466462

**Authors:** Alexander L. Cope, Premal Shah

## Abstract

Patterns of non-uniform usage of synonymous codons (codon bias) varies across genes in an organism and across species from all domains of life. The bias in codon usage is due to a combination of both non-adaptive (e.g. mutation biases) and adaptive (e.g. natural selection for translation efficiency/accuracy) evolutionary forces. Most population genetics models quantify the effects of mutation bias and selection on shaping codon usage patterns assuming a uniform mutation bias across the genome. However, mutation biases can vary both along and across chromosomes due to processes such as biased gene conversion, potentially obfuscating signals of translational selection. Moreover, estimates of variation in genomic mutation biases are often lacking for non-model organisms. Here, we combine an unsupervised learning method with a population genetics model of synonymous codon bias evolution to assess the impact of intragenomic variation in mutation bias on the strength and direction of natural selection on synonymous codon usage across 49 Saccharomycotina budding yeasts. We find that in the absence of *a priori* information, unsupervised learning approaches can be used to identify regions evolving under different mutation biases. We find that the impact of intragenomic variation in mutation bias varies widely, even among closely-related species. We show that the overall strength and direction of selection on codon usage can be underestimated by failing to account for intragenomic variation in mutation biases. Interestingly, genes falling into clusters identified by machine learning are also often physically clustered across chromosomes, consistent with processes such as biased gene conversion. Our results indicate the need for more nuanced models of sequence evolution that systematically incorporate the effects of variable mutation biases on codon frequencies.

## Introduction

Patterns of nucleotide base composition vary widely across a genome and are typically thought to be shaped by the interplay between adaptive and non-adaptive evolutionary processes. One common pattern observed in genomes across all domains of life is codon usage bias (CUB), the non-uniform usage of synonymous codons in coding sequences of genes [1, 2, 3, 4, 5]. Although synonymous nucleotide changes are often treated as neutral (i.e. codon frequencies are determined primarily by mutation bias and genetic drift) various lines of evidence indicate that synonymous changes are also subject to natural selection [6, 7, 8, 9, 10, 11, 12]. The coevolution between codon frequencies and tRNA pool, as well as the bias towards translationally efficient codons in highly expressed genes suggests translational selection is a major factor shaping genome-wide codon patterns [8, 13, 9, 14, 15, 16, 17, 18, 19]. Other selective forces, including selection against missense error [10, 20], selection against ribosome drop-off [21, 22], and selection to avoid mRNA secondary structure near the translation initiation site [23, 24], are also hypothesized to shape adaptive CUB. Although evidence indicates strong purifying selection can act on synonymous changes [25, 12], codon usage is generally thought to be subject to weak selection (i.e. *N*_e_*s* ≪ 1) with genome-wide codon frequencies at selection-mutation-drift equilibrium [26, 27, 16].

Translational selection is expected to be strongest in highly-expressed genes. Based on this assumption, various approaches for identifying the translationally preferred codon and quantifying selection on codon usage rely on comparing coding frequencies in highly-expressed genes to the remaining genome or lowly-expressed genes [15, 17, 19]. In contrast, other approaches quantify selection via the changes in codon frequencies as a function of gene expression, assuming that codon frequencies are at selection-mutation-drift equilibrium [16, 28, 29]. Failing to account for mutational biases can weaken or completely obfuscate signals of selection on codons usage, resulting in the development of various methods for separating out the effects of selection from mutation bias [30, 31, 32, 33, 34, 16, 29].

As translationsal selection is expected to produce a correlation between CUB and gene expression, a weak correlation between codon usage and gene expression could indicate low translational selection. Although current approaches often account for background mutational biases when estimating selection on codon usage, these models often assume a constant mutational bias across the genome. However, various processes cause the direction and strength of mutation biases to vary within the genome, potentially weakening signals of translational selection [35, 17, 36, 19]. Mutation rates can be context-dependent and vary depending on the identity of adjacent nucleotides in bacteria [37], yeast [38], and primates and humans [39, 40]. Isochores – long stretches of DNA with relatively homogeneous nucleotide composition – have been observed in eukaryotic species ranging from yeasts to humans [41, 42]. Various hypotheses propose to explain the presence of isochores, including variation in mutation biases related to the timing of genome replication and biased gene conversion [42, 43, 44]. Gene conversion – the transfer of genetic information from intact homologous sequences to regions containing double-strand breaks [45] – has been observed to be GC-biased in various phylogenetically-diverse species, resulting in increased GC content in recombination hotspots [46]. Lateral gene transfer events, including introgressions, can result in genes with distinct CUB being incorporated into a genome [47, 48]. Finally, mutation biases vary between leading and lagging DNA strands in some prokaryotic species [49, 50], leading to different CUB dependent upon the strand of a coding-sequence [51]. We know relatively little about the impact of variable mutation biases on the relationship between codon usage and gene expression, and how this impacts estimates of translational selection.

Population genetics models of coding sequence evolution are powerful tools for understanding the evolution of CUB [34, 16, 28]. Here we will use Ribosomal Overhead Cost version of the Stochastic Evolutionary Model of Protein Production Rates (ROC-SEMPPR) to estimate codon-specific estimates of natural selection and mutation bias, as well gene-specific estimates of protein production rates [29]. Recent work applied ROC-SEMPPR to quantify the differences in codon usage between the ancestral genome and a large introgression in *Lachancea kluyveri*, finding that the ability to detect selection on codon usage improved if assuming codon usage in the introgressed region was shaped by different mutational and selective biases than the ancestral genes [48]. Here, we use ROC-SEMPPR to examine the effects of within-genome variation in mutation biases on the ability to detect natural selection on codon usage across the Saccharomycotina budding yeast subphylum [18]. Unlike *L. kluyveri, a priori* knowledge of genes evolving under different mutational biases is lacking in a large number of these species. To hypothesize genes shaped by different mutational biases, we apply an unsupervised machine-learning approach based on codon frequencies previously described in [52]. We highlight various yeasts in which intragenomic variation in mutational biases masks the efficacy of natural selection on codon usage. We find that genes falling into the same clusters determined via the unsupervised machine-learning algorithm are physically clustered within the genome, suggestive of processes such as biased gene conversion.

## Results

### Relationship between observed and predicted gene expression varies significantly between closely-related species

ROC-SEMPPR was applied to the nuclear coding genes of 49 species from the Saccharomycotina budding yeast subphylum for which we were able to obtain empirical estimates of mRNA abundances (see Methods). We will refer to these fits as the constant mutation (“ConstMut”) model as mutation bias parameters are assumed to be the same across all coding sequences. We use closely-related species across three genera (*Saccharomyces, Candida*, and *Ogataea*) to highlight the differences in relationship between empirical gene-expression levels and expression levels predicted based on codon usage patterns using a stochastic evolutionary model (ROC-SEMPPR). Across all three genera, one of the species shows a relatively high correlation between empirical and predicted expression levels, while the other species show a weak or even negative relationship (Figure 1A). These discrepancies might be due to three reasons - (i) poor quality of expression data for these non-model organisms, (ii) rapid changes in degree of translation selection across closely related species, or (iii) inaccurate predictions of gene-expression data from CUB due to model mis-specification.

**Figure 1:**
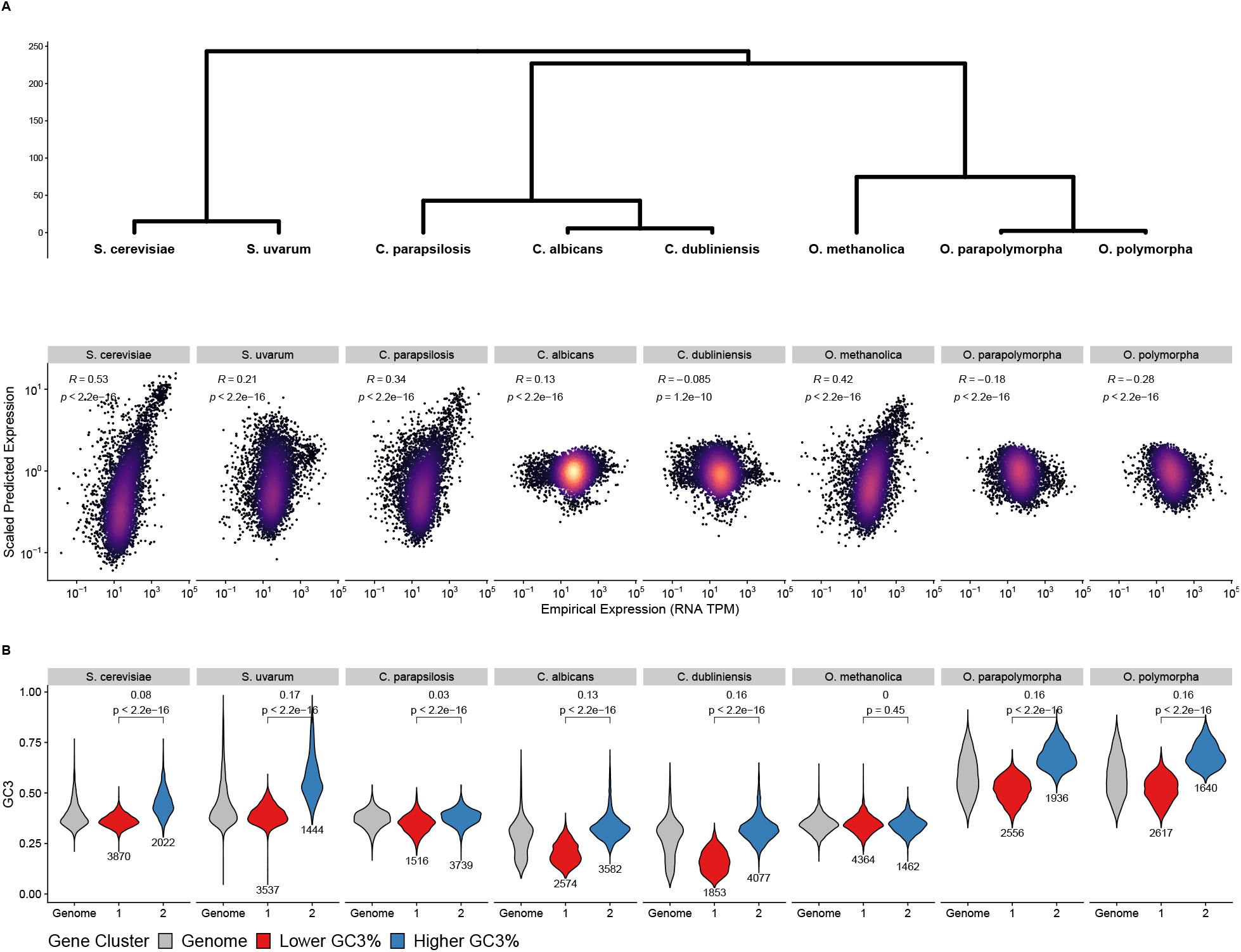
(A) Correlations between empirical mRNA abundances and predicted expression values from ROC-SEMPPR across 8 Saccharomycotina species. Phylogenetic relationships between species indicates relatedness between species, with branch lengths units in millions of years. (B) Distribution of GC3% for the entire genome (grey), as well as for the distributions within each of the clusters determined by the CLARA clustering. We denote the cluster with the lower median GC3% as the Lower GC3% cluster (red) and the cluster with the higher median GC3% as the Higher GC3% cluster (blue).Brackets indicate the difference in medians between the lower and higher GC3% clusters, as well as the p-value from a Kolmogorov-Smirnov test comparing the distribution of GC3% between the two clusters.

When comparing expression data between closely-related species we find that they are of similar sequencing depth and are moderately correlated with each other (Supplementary Figure 1A). Taken together, the evidence indicates that variation in quality of gene-expression datasets is insufficient to explain these large changes in correlation between observed and predicted expression levels across closely-related species.

To examine if the degree and direction of translation selection differs between closely-related species of the three genera, we compared the relative synonymous codon usage (RSCU) between genes with high and low expression levels (top 5% and bottom 5%, respectively). Across all three genera, RSCU values calculated from highly-expressed genes are well-correlated, indicating that the overall direction of selection on synonymous codon usage is similar between species (Supplementary Figure 2). For all 8 species, RSCU values from low and high expression genes are correlated, but the RSCU values of many codons clearly differ between high and low expression genes (Supplementary Figure 3). Taken together, these results suggests the strength and direction of selection on synonymous codons is similar across species within the same genus.

### Intragenomic variation in mutational biases affects estimates of translational selection

Models of sequence evolution of codon usage typically assume that the mutational biases associated with a particular synonymous codon are constant across the entire genome. However, several non-adaptive forces such as biased gene-conversion [46], context-dependent mutation rates [37], and distribution of genes across different isochoric regions [42, 43] in the genome can affect the local mutation rates and biases and therefore affect codon composition. To account for this local variation in mutational biases, we let genes evolve under one of two mutational regimes but under shared selection constraints. Genes were assigned to mutational regimes using a k-medoids CLARA clustering algorithm applied to the principal axes estimated from absolute codon frequencies of each species (see Methods). The two clusters differed in terms of average GC3% content (Figure 1B), suggesting some genes (designated as the Higher GC3% cluster) may be subject to stronger GC mutational biases. A clear pattern emerges when examining the differences in GC3% content between the Lower GC3% and Higher GC3% cluster: species with larger differences (i.e. > 0.1) in the median GC3% content between clusters also have weaker or negative correlations between predicted and empirical gene expression (Figure 1B,C). We note that the distributions of GC3% between clusters for each species, with the exception of *O. methanolica*, are significantly different (Kolmogorov-Smirnov test, *p* < 2.2*e −* 16).

ROC-SEMPPR was refit to all species allowing for mutation bias parameters to vary across the two subsets of coding sequences determined by the clustering algorithm, although selection parameters were assumed to be the same due to these genes evolving under the same tRNA pool. We will refer to these model fits as the varying mutation (“VarMut”) model. Species showing greater differences in GC3% between the clusters showed significant improvement in the correlation between predicted and empirical gene expression when accounting for intragenomic variation in mutation bias (Figure 2A,B). For species with relatively little difference in GC3% between clusters, the VarMut model often resulted in a poorer agreement between empirical and predicted expression estimates. This suggests that clustering in cases where intragenomic variation in mutation bias is small or non-existent is diluting signals of selection on codon usage. In the 5 species for which the VarMut model fit significantly improved predictions of gene expression, the predicted expression estimates from the ConstMut model fit essentially captures the Lower GC3% and Higher GC3% clusters (Figure 2C). This indicates that variation in mutational biases, when strong enough, can be mistaken for signals of natural selection.

**Figure 2:**
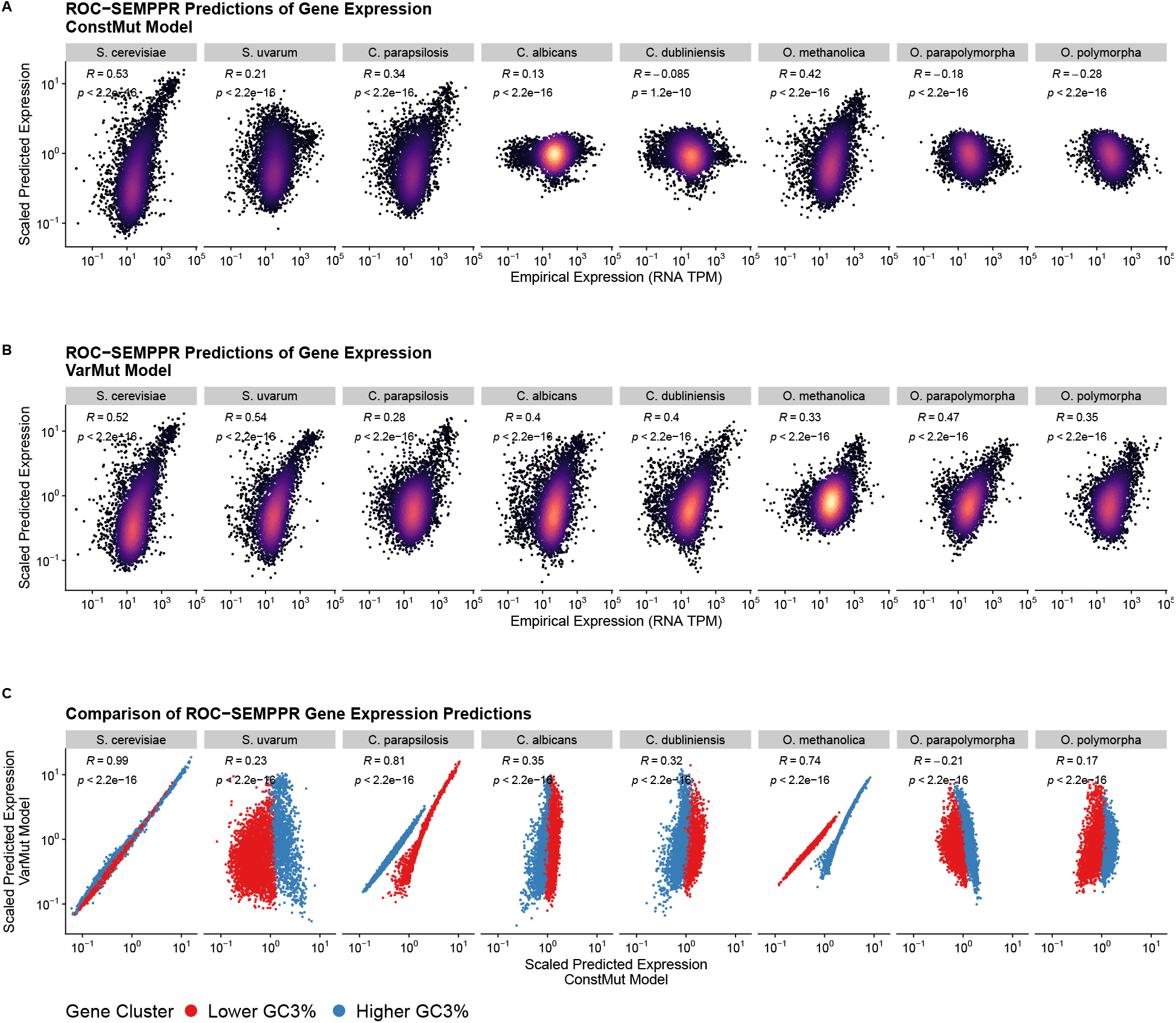
Effects of allowing for mutation bias variation on predictions of gene expression. (A) Correlation between empirical and predicted gene expression values when mutation bias parameters are shared across clusters (ConstMut Model), as in Figure 1. (B) Correlation between empirical and predicted gene expression values when mutation bias parameters are allowed to vary across clusters (VarMut model). (C) Comparison of predicted gene expression between the ConstMut and VarMut models.

### Genes in a mutational cluster are physically clustered on chromosomes

Results clearly indicate that differences in mutational biases across genes mask signals of selection on codon usage in species showing greater variability in GC3%. To understand which molecular processes might be responsible for the clustering of genes in different GC3% categories, we checked if genes belonging to individual clusters are in close proximity to each other on the chromosome. Presence of large physical clusters of genes belonging to each mutational clusters could indicate the role of biased gene conversion in affecting differential mutational rates along a chromosome. Failure to observe such physical clustering on chromosomes of high or low GC3% genes would suggest novel mechanisms of variation in mutation biases. To quantify the degree of physical clustering of genes belonging to each mutational cluster, we estimated GC3% variation along chromosomes using either a 20-gene moving average approach or 20-gene non-overlapping windows.

We observe that genes assigned to the same mutational cluster are also physically clustered across the chromosomes of species for which the VarMut model was better able to detect selection on codon usage relative to the ConstMut model (Figure 3 and Supplementary Figures 4 – 11). As with the mutational clusters (Figure 1C), physical clusters of genes assigned to the same mutational cluster reflect relatively large (sometimes spanning hundreds of genes) GC3-rich or GC3-poor regions (Figure 3, left panels). Using non-overlapping 20-gene windows, which allows us to apply statistics that assume independent data, we observe that the average GC3% within a region is highly-correlated with the percentage of genes assigned to the Higher GC3% within that region across all chromosomes of a species (Figure 3, right panels). However, physical clustering of genes assigned to the same mutational cluster was overall weaker across species for which the VarMut model failed to improve ability to detect selection on codon usage (Supplementary Figure 12).

**Figure 3:**
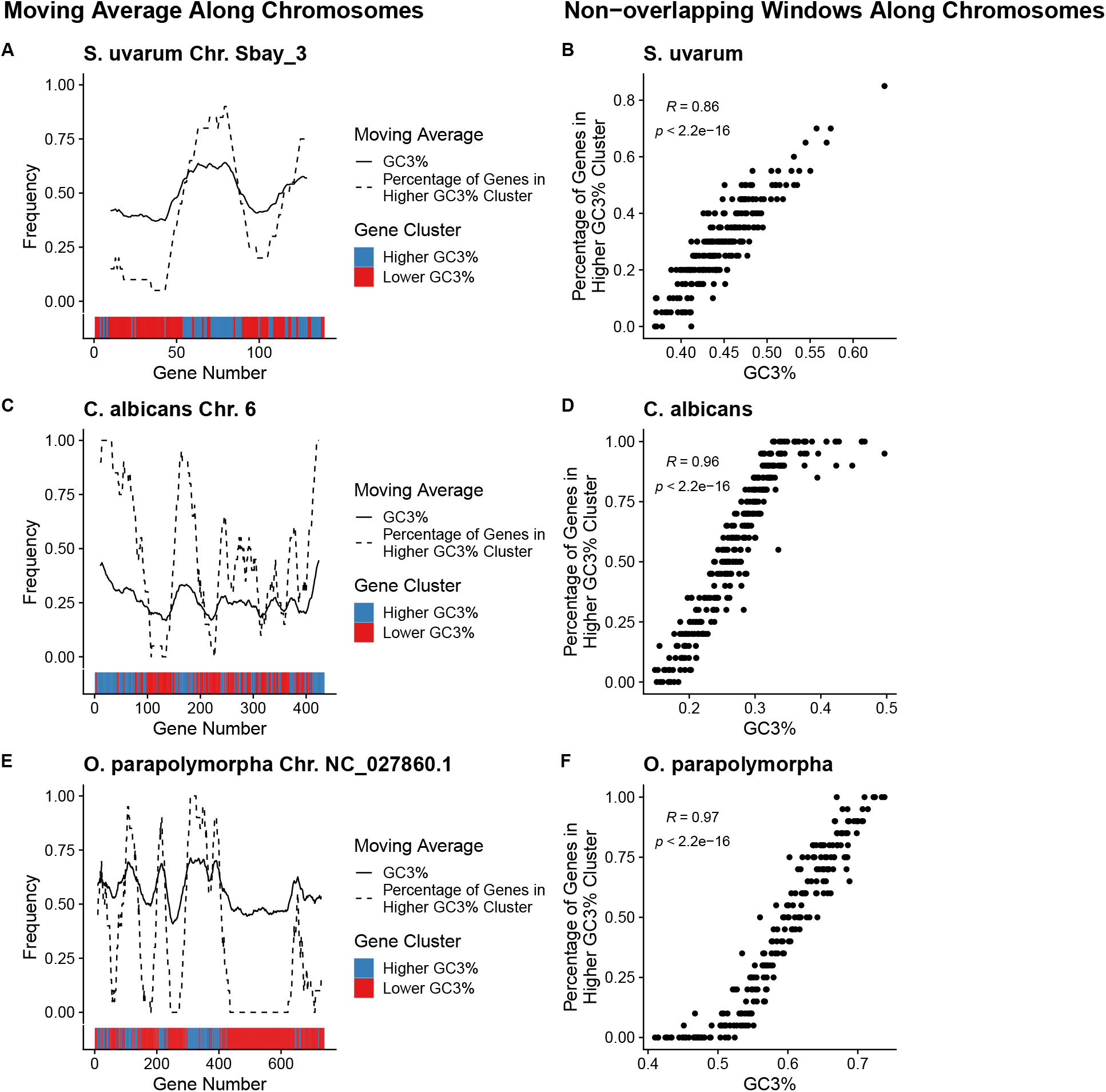
(Left panel) Per-gene GC3% content across select chromosomes quantified as a moving average using a 20 gene sliding window (solid line). For each 20 gene window, the percentage of genes assigned to the Higher GC3% regime is also shown (dashed line). Color bars indicate the mutation regimes for Higher and Lower GC3% (blue and right, respectively). (Right panel) Scatter plots showing per-gene GC3% and percentage of genes assigned to the Higher GC3% regime using non-overlapping 20 gene windows across all chromosomes. The independence of windows allows us to apply the Spearman Rank correlation. (A,B) *S. uvarum*. (C,D) *C. albicans*. (E,F) *O. parapolymorpha*.

The physical clustering of genes with high and low GC3% is consistent with GC-biased gene conversion (gBGC) [46]. gBGC occurs in the *sensu stricto* yeasts; however, [17] concluded it has the strongest impact on the shaping the CUB of *S. uvarum*, consistent with our results. Unlike with *Saccharomyces, C. albicans* and *C. dubliniensis* do not have a sexual cycle, eliminating meiotic recombination as a mechanistic basis. However, [44] proposed mitotic recombination as potential mechanism for gBGC.

### Effects of Variable Mutation on Estimates of Natural Selection

Allowing for intragenomic variation in mutation model improves detection of natural selection on codon usage. A key aspect to studying CUB is identifying the “optimal” or “preferred” codon. Across the 5 species for which we found the VarMut model outperformed the ConstMut model, the optimal codon was misidentified for the majority of amino acids by latter (Figure 4B). In the case of amino acids for which the optimal codon was the same between models, selection coefficient estimates for many codons showed large changes between the models (Figure 4C). This indicates that the strength of selection may be over or under-estimated when intragenomic mutation bias variation is unaccounted for, even if the optimal codon is correctly identified.

**Figure 4:**
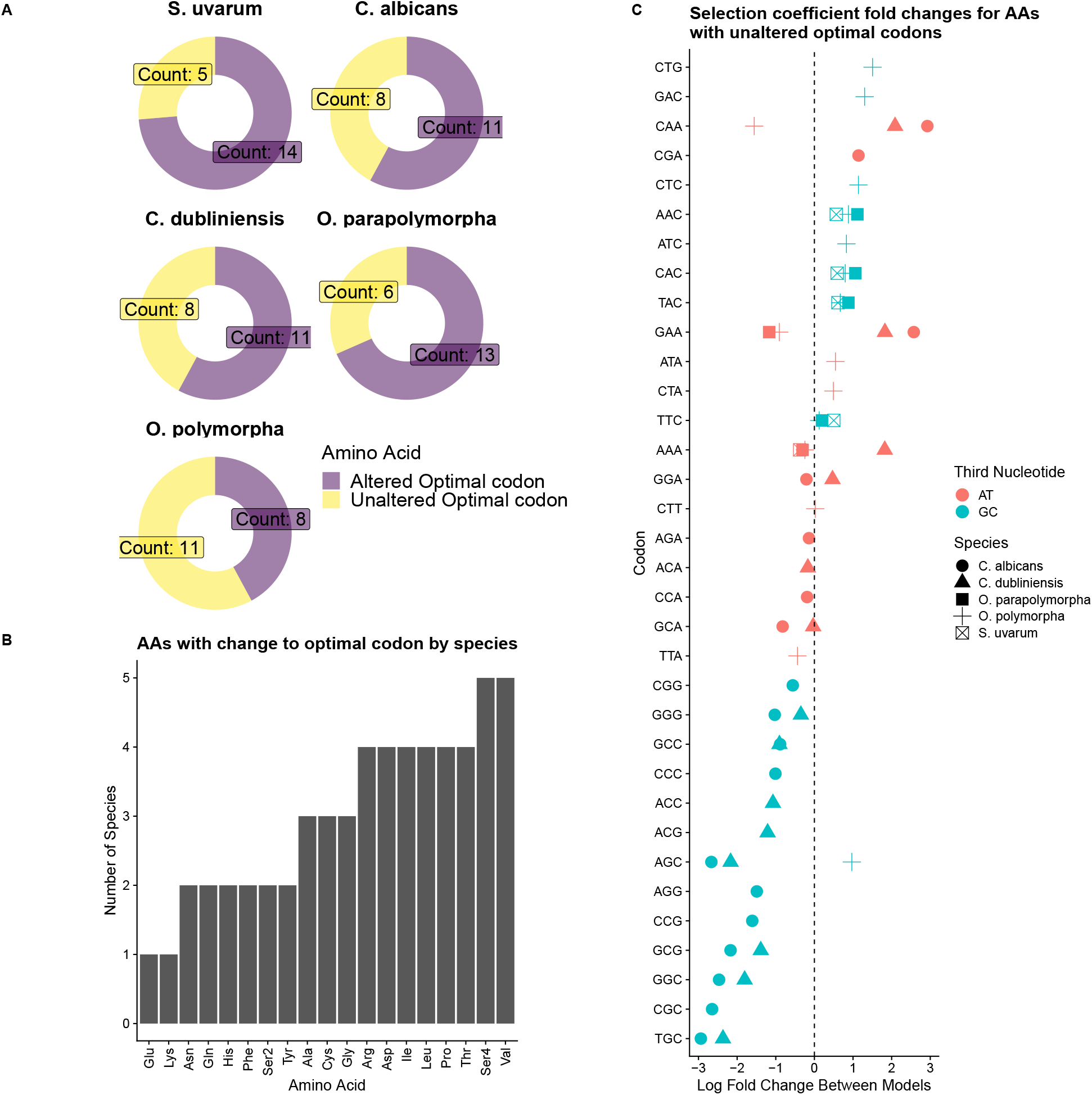
Data represent results from *S. uvarum, C. albicans, C. dubliniensis, O. parapolymorpha*, and *O. polymorpha*. (A) Number of amino acids for which the optimal codon differs between the ConstMut and VarMut models. (B) Breakdown of fold changes for amino acids in which optimal codon is the same between ConstMut and VarMut models.

Selection coefficient estimates from ROC-SEMPPR represent the preference of a codon relative to a single reference codon. As the clusters reflect differences in GC3%, selection coefficients were rescaled to represent selection in NNG vs. NNA or NNC vs. NNT ending codons. GC-ending codons are universally disfavored by natural selection relative to AT-ending codons in *C. albicans* when fitting the ConstMut model (Supplementary Figure 13A). This is consistent with the distributions of predicted expression estimates, with genes falling into the Higher GC3% cluster predicted to have lower expression (Figure 2C). The VarMut model reveals that some GC-ending codons are favored by natural selection relative to the corresponding AT-ending codon (Supplementary Figure 13A, bottom right quadrant). Even in cases where the GC-ending codon remains disfavored relative to the AT-ending codon, we find that the magnitude of the estimated selection coefficients is lower in the VarMut model (Supplementary Figure 13A), indicating selection against these codons is weaker than suggested by the ConstMut model. Other species for which the VarMut model improved ROC-SEMPPR’s ability to detect natural selection on codon usage showed similar results (Supplementary Figure 13B). Unsurprisingly, we find that mutation bias estimates against GC-ending codons (again, relative to AT-ending codons) is much weaker in the Higher GC3% cluster than the Lower GC3% cluster in these species (Supplementary Figure 14).

### Detection of selection on CUB improves when allowing for intragenomic mutation bias variation in species across the Saccharomycotina subphylum

Up to this point, results have primarily focused on 8 species represented by three genera within the Saccharomycontina subphylum. To understand the effects of intragenomic mutation bias variation across the larger phylogeny, we expanded our analysis to 49 species across the subphylum for which reliable expression datasets are available (Figure 5A). Accounting for intragenomic mutation bias variation has a relatively small impact on the detecting selection on codon usage for the majority of species (Figure 5A). However, the correlations between empirical and predicted genes expression estimates improves in various species, indicating that intragenomic variation in mutation biases is present across the Saccharomycontina subphylum. We find that species with larger variation in GC3% content, quantified as the difference in median GC3% content between the Lower and Higher GC3% clusters, tend to show larger improvements in the correlations between empirical and predicted expression estimates when fitting the VarMut Model (Spearman Rank correlation *r* = 0.44,*p* = 0.0019, Figure 5B). This correlation remains statistically significant after transforming values using phylogenetic independent contrasts (Spearman Rank Correlation *r* = 0.48, *p* = 0.0006) [53].

**Figure 5:**
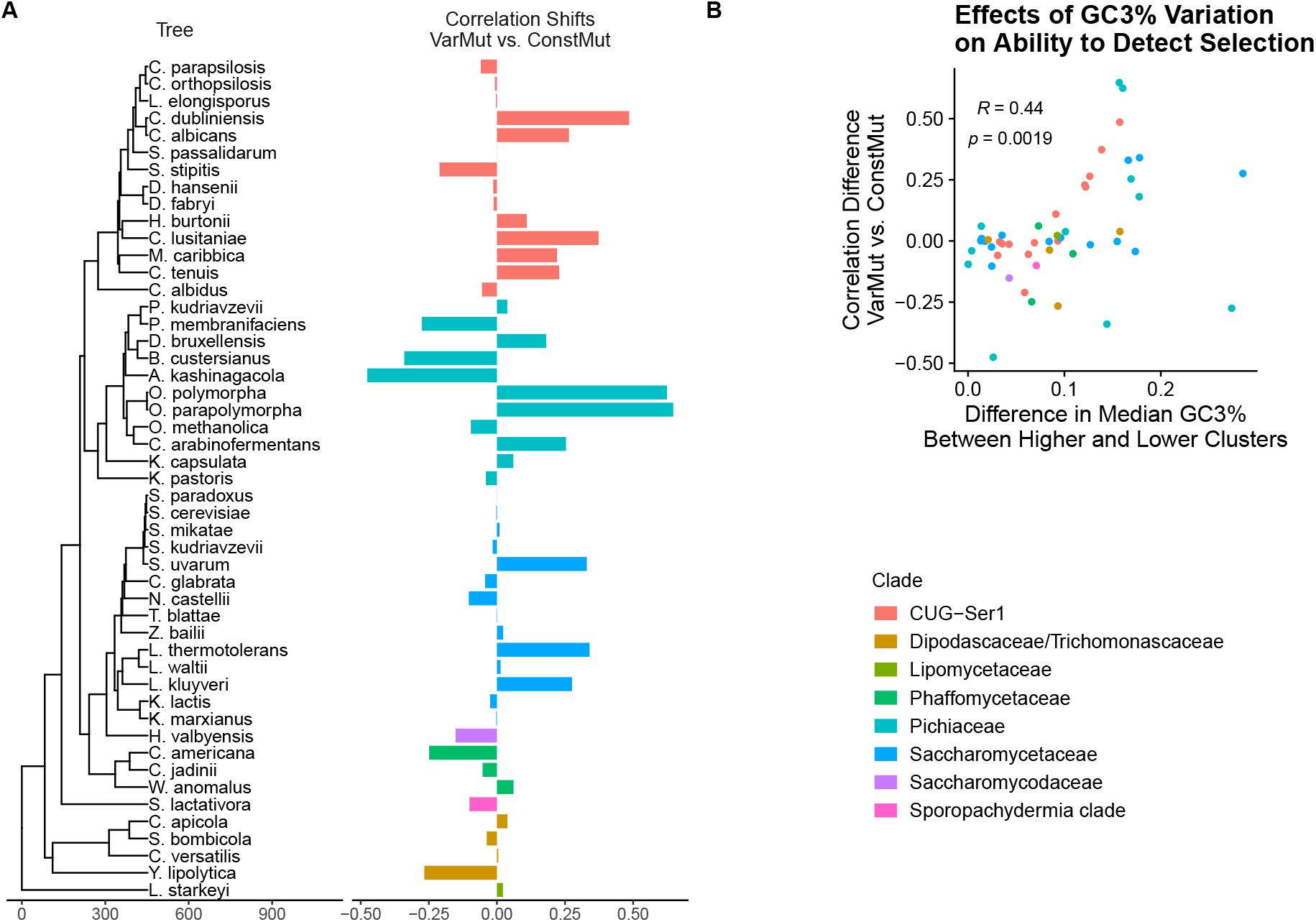
Clade names are taken from [18]. (A) Change in Spearman correlation between empirical and predicted gene expression when fitting VarMut model relative to ConstMut model. A positive value indicates an improvement in correlation when fitting the VarMut model. (B) Relationship between the shift in correlations as a function of the difference in median GC3% content between the Lower and Higher GC3% clusters.

Interestingly, the effect of intragenomic variation in mutation bias can vary between closely related species, as previously highlighted in species from the *Saccharomyces, Candida*, and *Ogataea* genera. In addition, the causes of intragenomic variation in mutation bias can also vary between closely related species. For example, both *L. kluyveri* and *L. thermotolerans* show a substantial improvement in the predictions of gene expression with the VarMut model; however, only the former contains the introgressed region on one of its chromosomes [48]. In the case of *L. kluyveri*, the Higher GC3% cluster is predominantly the introgressed region (Supplementary Figure 15). In contrast, *L. thermotolerans* shows relatively large variation in GC3% content along its chromosomes, similar to *C. albicans* and *C. dubliniensis* (Supplementary Figure 16).

## Discussion

A key to understanding the evolution of codon usage bias (CUB) across a phylogeny is to quantify the relative contributions of non-adaptive (e.g. mutation bias) and adaptive evolutionary forces in shaping codon frequencies. Studies often assume that mutation bias is constant across the genome; however, this may not always be the case due to processes such as biased gene conversion (BGC). Across 49 Saccharomycontina yeasts, failing to account for intragenomic variation in mutation bias obfuscates signals of natural selection on codon usage in various phylogenetically-diverse species. This includes misidentification of the direction of selection (i.e. misidentification of optimal codons), and the over/under-estimation of the strength of selection. In the absence of *a priori* hypotheses, such as large introgression events [48], unsupervised clustering of genes based on codon frequencies can reveal genes evolving under different mutational biases. Interestingly, we observe in many species that the unsupervised clustering is mirrored by physical clustering along chromosomes, i.e. genes falling into the same mutational cluster tend to be physically grouped together within the genome. For some species, such as *C. albicans* and *O. parapolymorpha*, this is a constant across chromosomes, with each chromosome shows large variations in GC3% content. For other species, such as *S. uvaraum*, regions of large variation in GC3% content appear to be more isolated. Species showing little variation in GC3% content across their chromosomes showed little, if any improvement, by allowing for intragenomic variation in mutation bias.

To categorize genes evolving under different mutation biases, we used the CLARA clustering algorithm (a type of k-medioids clustering) and specifically clustered genes into two groups (*k* = 2). It is possible that identifying more clusters results in further improvements in detection of natural selection on codon usage. However, it is important to remember that for most species under strong translational selection, a major factor shaping variation in codon frequencies is gene expression. Over-clustering of genes could potentially dilute out the signals of natural selection by misidentifying genes with high expression and altered codon frequencies as evolving under a different mutational regime. We find evidence of this in species that have low variation in GC3% along the genome, such as *C. parapsilosis*, in which the VarMut model performs significantly worse at predicting gene expression.

The physical clustering of genes along a chromosome that are assigned to either the Higher or Lower GC3% mutation regimes, is consistent with the action of GC-biased gene conversion (gBGC). gBGC has been observed in *S. cerevisiae* [54], but its overall impact on codon frequencies has been a point of debate [55, 56, 57, 58, 17]. We observed that the VarMut model had a negligible impact on our ability to detect natural selection in *S. cerevisiae* and other *sensu stricto* yeasts. Despite showing little variation in GC3% along its chromosomes compared to species from the the *Candida* and *Ogataea* genera, intragenomic variation in mutation biases appear to be strong enough to obscure signals of selection in *S. uvarum*, but no other *sensu stricto* yeasts. Surprisingly, ROC-SEMPPR occasionally encountered a local maximum when fit to *S. uvarum* that appeared to adequately capture the effects of selection on codon usage. This is consistent with previous work that concluded gBGC had the largest impact on codon usage in *S. uvarum* among the *sensu stricto* yeasts, although the effect was weak relative to selection across all species [17].

Although *C. albicans* and *C. dubliniensis* both show physical clustering on the chromosomes of GC3 rich and GC3 poor genes, these species are not known to reproduce sexually, eliminating meiotic gene conversion as a possible source of gBGC. However, both *C. albicans* and *C. dubliniensis* can reproduce via a “parasexual” or “parameiosis” cycle [59, 60]. Previous analysis has shown that mitotic gBGC could cause the observed variation in GC3% [44]. Interestingly, a recent analysis observed that interlocus or nonallelic gene conversion, in which paralogous sequences are used to repair double-strand breaks, is GC-biased in the the pathogenic yeast *Candida krusei* – the anamorph of *Pichia kudriavzevii* [61, 62]. However, *P. kudriavzevii* showed little improvement when fit by the VarMut model. Further work is needed to understand the underlying causes of intragenomic variation in mutation biases across the Saccharomycotina species.

Tests of selection on codon usage are often based on comparing the codon usage of a gene to a reference set thought to be biased towards selectively-favored codons. These tests often create a reference set using assumed highly expressed genes (e.g. ribosomal proteins [63], but see [64]) or using empirical gene expression data [19]. In the case of intragenomic variation in mutation bias, the reference set must be chosen to minimize the impact of the variable mutation biases, as this may result in an over or underestimation of the strength of selection. Even if care is taken in choosing the reference set, variable mutation biases may still lead to the over or underestimation of selection on codon usage in specific genes. For example, if selection generally favors GC-ending codons, then natural selection on codon usage may be overestimated in low expression genes subject to gBGC.

Approaches for dealing with variation in mutation biases rely on the analysis of SNPs detected across multiple individuals from a population, or comparisons of codon and nucleotide frequencies in regions of low and high recombination [17, 35, 65, 66, 36]. However, polymorphism and recombination data are often unavailable for non-model species, limiting the applicability of these approaches. We combined unsupervised machine-learning with an explicit populations genetics model to estimate selection on codon usage in the context of variable mutational biases. Although machine-learning is a powerful tool, the descriptive nature of such approaches can make biological interpretations difficult. As our understanding of the coding sequence evolution matures, models that more explicitly incorporate the various evolutionary forces that shape codon usage patterns, such as differences in mutation bias due to gBGC, are necessary.

## Materials and Methods

We obtained genome sequences and associated annotation files from [67]. We excluded mitochondrial genes, protein-coding genes with non-canonical start codons, internal stop codons, and sequences whose lengths were not a multiple of three from all analysis. To identify mitochondrial genes, we queried all protein sequences against a BLAST database built from sequences in the MiToFun database (http://mitofun.biol.uoa.gr/).

Empirical mRNA abundances were used as a proxy of protein production rates. For each species, publicly available RNA-seq data were downloaded from the Sequence Read Archive or the European Nucleotide Archive (*L. kluyveri only*). Adapters for each sequence were trimmed using fastp [68] and genes were quantified using kallisto [69]. Transcripts-per-million (TPMs) were re-calculated for each transcript by rounding raw read counts to the nearest whole-number [70].

### Identifying Intragenomic Variation in Codon Usage Bias

To identify genes potentially evolving under different mutational biases, an unsupervised learning approach was implemented as described in [52]. Correspondence Analysis (CA) was performed for each species based on absolute codon frequencies using the ca R package [71]. We note that Relative Synonymous Codon Usage (RSCU) was not used as it has been found to introduce artifacts to CA [72]. Genes were then clustered into two groups based on the first 4 principle components from the CA using the CLARA algorithm implemented in the cluster R package[73]. The CLARA algorithm is designed to perform k-medoids clustering on large datasets. For each species, the cluster with the lower median GC3% was designated as “Lower GC3% Cluster” and the cluster with the higher median GC3% was designated as the “Higher GC3% Cluster”.

### Analysis of CUB

ROC-SEMPPR was fit to 49 species using the R package AnaCoDa [74]. ROC-SEMPPR is implemented in a Bayesian framework, allowing it to simultaneously estimate codon-specific estimates of selection coefficients and mutation bias with gene-specific estimates of the evolutionary average gene expression by assuming gene expression is lognormally distributed [29]. This allows the model to be fit to any species with annotated protein-coding sequences. Parameters are estimated via a Markov Chain Monte Carlo (MCMC). For each species, MCMC were fit for 200,000 iterations, keeping every 10^th^ iteration. The first 50,000 iterations were discarded as burn-in. For each analysis, two separate chains were run from different starting points and parameter estimates were compared to assess convergence.

To handle species for which CTG codes for serine instead of the canonical amino acid leucine, a local version of AnaCoDa was created that is capable of handling this amino acid switch. As ROC-SEMPPR assumes weak mutation (i.e. *N*_e_*µ ≪* 1), CTG was also excluded from the set of serine codons [29]. For each species, ROC-SEMPPR was first fit assuming selection coefficient and mutation bias parameters were the same between the two clusters, which we refer to as the “ConstMut” model. Each was species was then fit allowing the mutation bias to vary for genes indicated by the two clusters, which we will refer to as the “VarMut” model. For the VarMut model, selection coefficients were assumed to be the same across clusters as codon usage in both clusters is still adapting to the same tRNA pool [75].

ROC-SEMPPR predictions of gene dexpression were compared to empirical estimates of mRNA abundance using the Spearman rank correlation coefficient. If natural selection on codon usage is strong enough to shape codon usage frequencies, then a statistically-significant positive correlation is expected between predicted and empirical estimates of gene expression [76, 34, 29]. If mutation bias varies across the genome, then it is expected that allowing this term to vary between clusters will significantly improve the correlation between predicted and empirical gene expression. To assess the impact of evolutionary processes that may favor GC-ending codons over AT-ending codons, such as gBGC, selection coefficient estimates and mutation bias estimates were rescaled such that all GC-ending codons were relative to the corresponding AT-ending codon, e.g. GCG was scaled relative to GCA. In this context, a positive selection coefficient (mutation bias) estimate indicates the GC-ending codon is disfavored by natural selection (mutation bias) relative to the AT-ending codon. Similarly, a negative value indicates the GC-ending codon is favored relative to the AT-ending codon.

### Examining variation in GC3% along chromosomes

Genes were mapped to their corresponding location on each chromosome indicated within the genome. Genes filtered out prior to analysis with ROC-SEMPPR were also excluded from this analysis. Variation in GC3% content along chromosome was visualized using a moving average of the per-gene GC3% value across a 20 gene window. Along with this, we calculated the percentage of genes falling into the Higher GC3% cluster within the same 20 gene window. This provides a means to compare GC3% variation along a chromosome with the cluster membership and provides insights into if genes falling into the two clusters determined by the CLARA clustering are also physically clustered within the genome. To deal with non-independence due to autocorrelation, we also calculated the same metrics using non-overlapping 20 gene windows, which were then compared using a Spearman Rank correlation.

### Comparisons across species

To visualize results across 49 species, we used the ggtree R package [77]. The phylogenetic tree was taken from the Supplementary Material of [67]. Phylogenetic independent contrasts (PIC) was used to account for the non-independence when calculating correlations across species [53].

## Data Availability

No new data were generated for this project. All data are publicly available via the citations provided in this article. Scripts for re-creating our analysis and visualizations can be found at https://github.com/acope3/Intragenomic_variation_mutation_bias.

## Acknowledgments

The authors would like to thank E.W.J. Wallace and M.A. Gilchrist for helpful discussions over the course of this project. A.L.C would like to thank the INSPIRE (IRACDA New Jersey/New York for Science Partnerships in Research and Education) Postdoctoral Program (NIH PAR-19-366) for current financial support. This work was supported by the National Science Foundation (DBI 1936046) and the National Institutes of Health (R35 GM124976) awarded to P.S.

## Supplementary Figures

**Figure 1:**
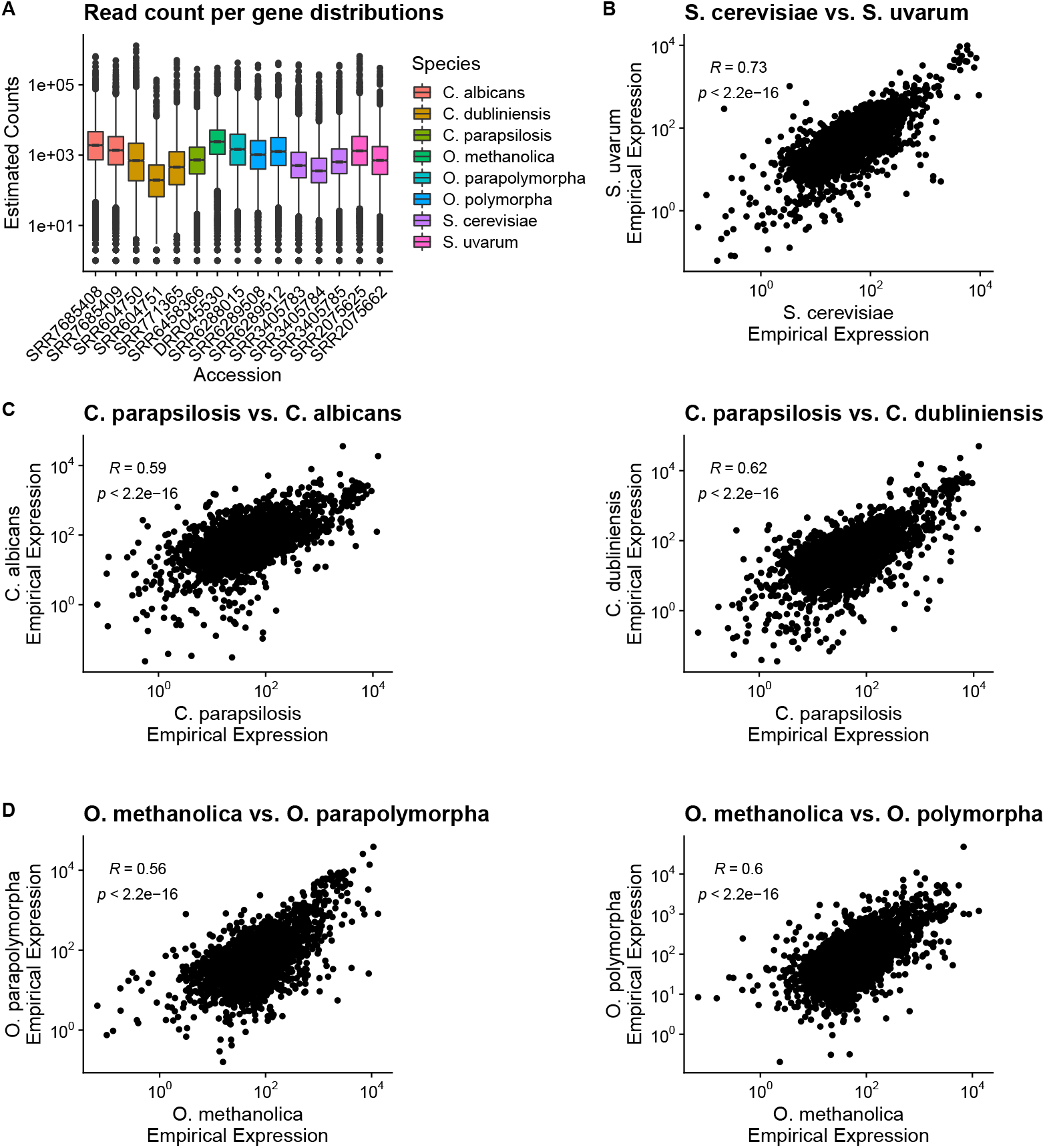
Comparison of empirical mRNA abundances estimated from disparate RNA-seq measurements. **(A)** Distribution of per-gene estimated counts from kallisto. **(B-D)** Comparison of empirical expression estimates across species using one-to-one orthologs. Correlations represent Spearman rank correlation coefficients.

**Figure 2:**
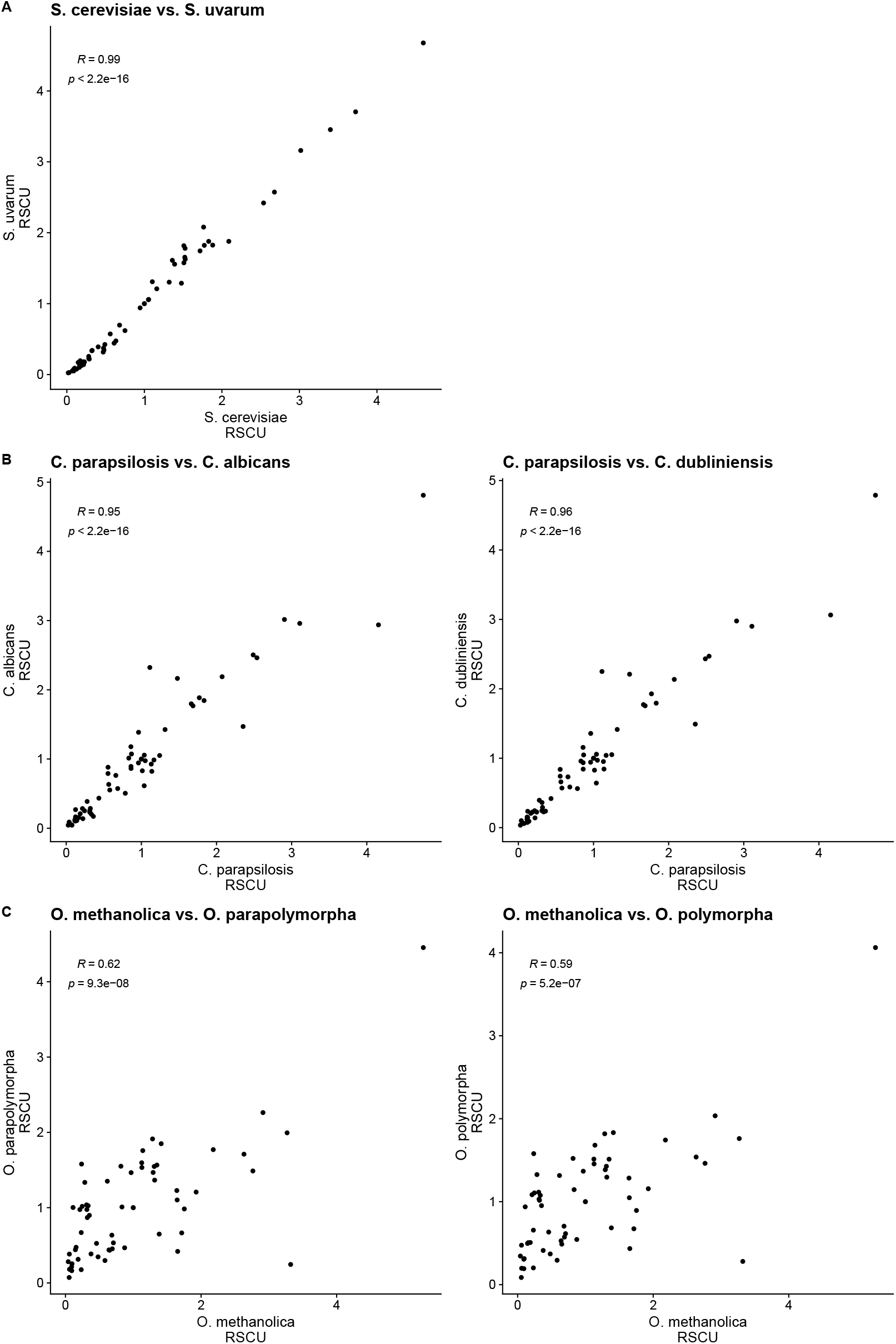
Across-species comparison of RSCU values calculated from the most highly expressed genes (top 5% based on empirical expression estimates). **(A)** *Saccharomyces*. **(B)** *Candida*. **(C)** *Ogataea*. Correlations represent Spearman rank correlation coefficients.

**Figure 3:**
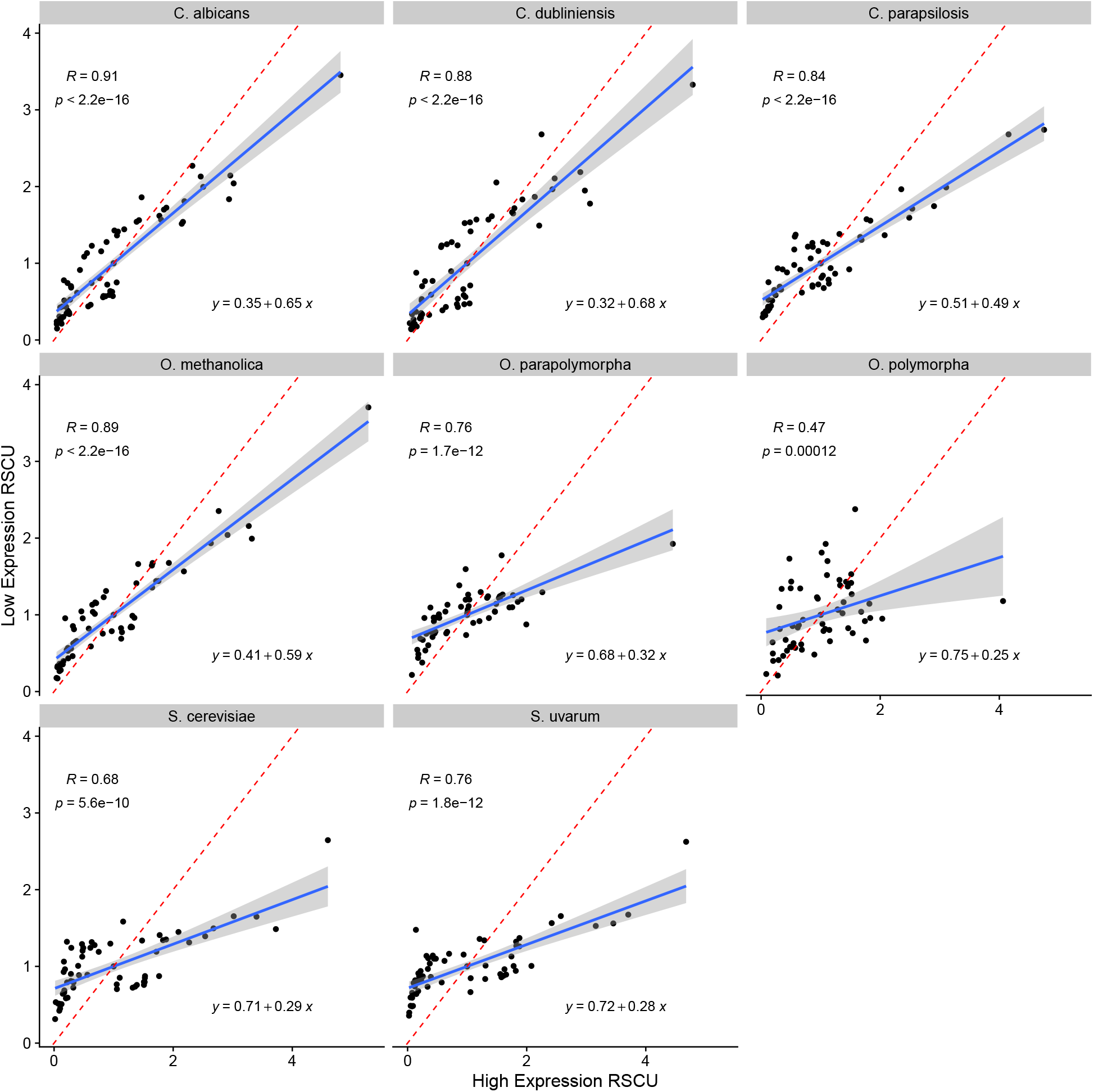
Comparison of RSCU values calculated from most highly and lowly expressed genes (top 5% and bottom 5% of expression estimates) for each species. Correlations represent Spearman rank correlation coefficients.

**Figure 4:**
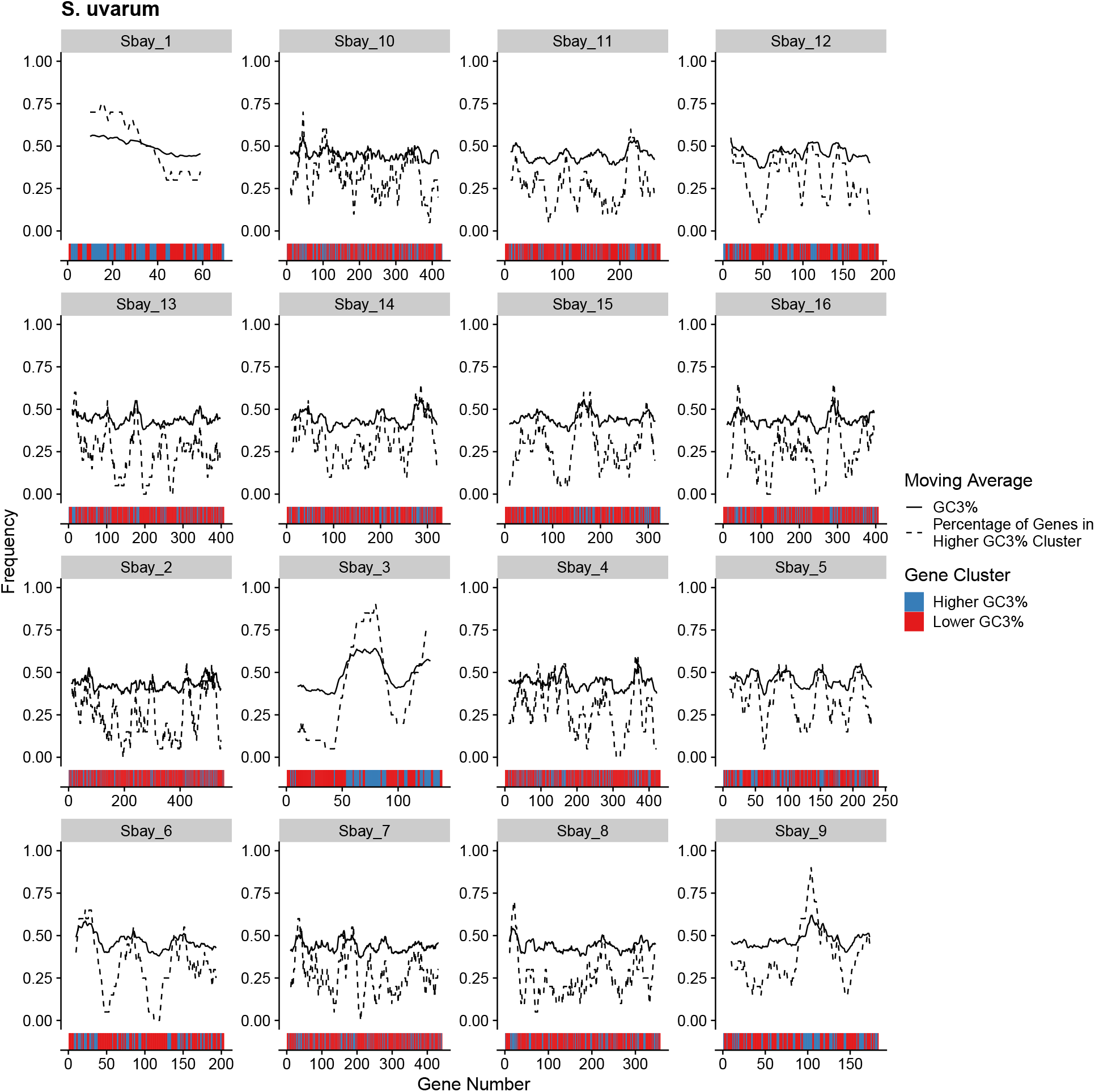
Per-gene GC3% content across all *S. uvarum* chromosomes quantified as a moving average using a 20 gene sliding window (solid line). For each 20 gene window, the percentage of genes assigned to the Higher GC3% regime is also shown (dashed line). Color bars indicate the mutation regimes for Higher and Lower GC3% (blue and right, respectively).

**Figure 5:**
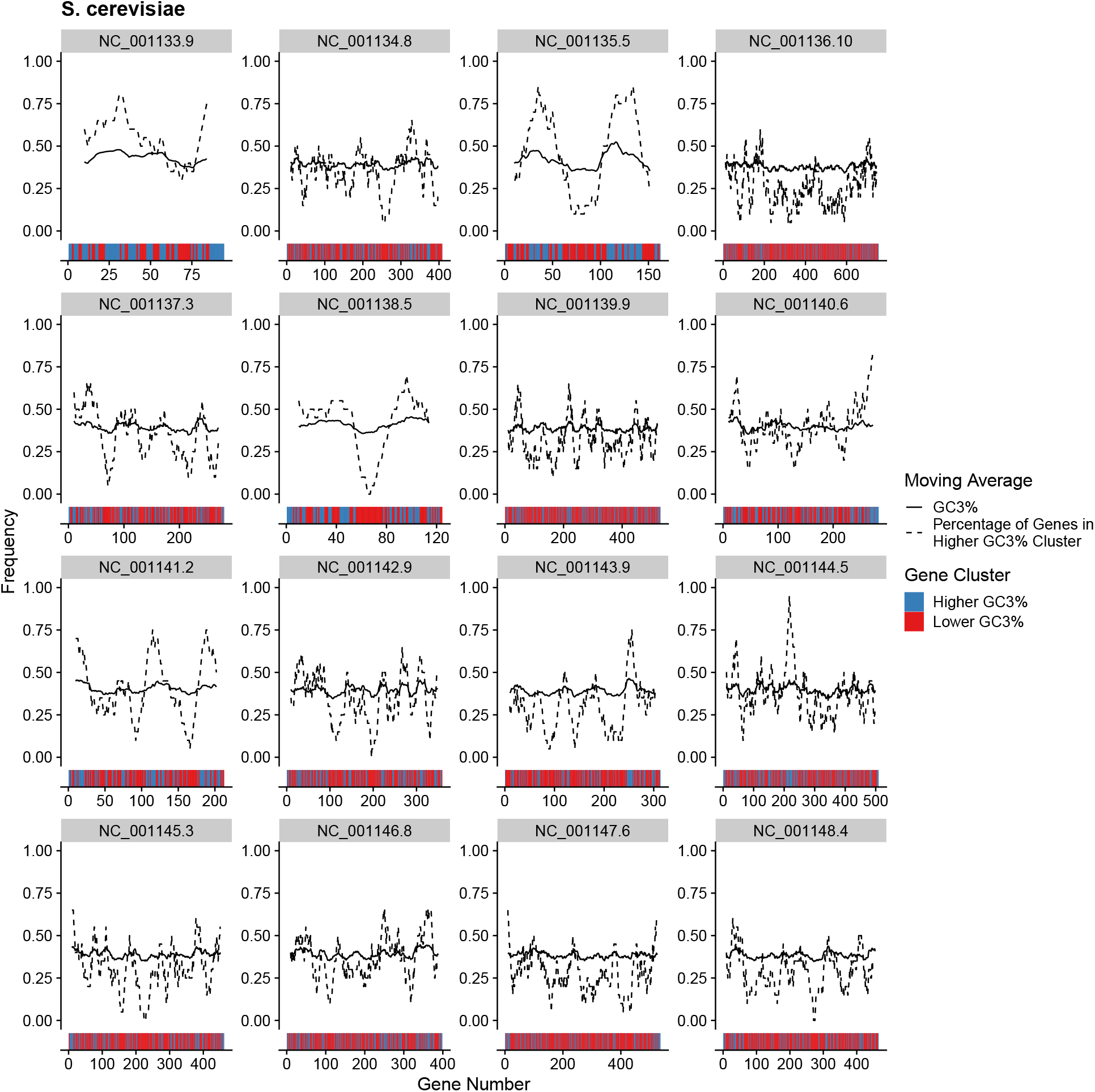
Per-gene GC3% content across all *S. cerevisiae* chromosomes quantified as a moving average using a 20 gene sliding window (solid line). For each 20 gene window, the percentage of genes assigned to the Higher GC3% regime is also shown (dashed line). Color bars indicate the mutation regimes for Higher and Lower GC3% (blue and right, respectively).

**Figure 6:**
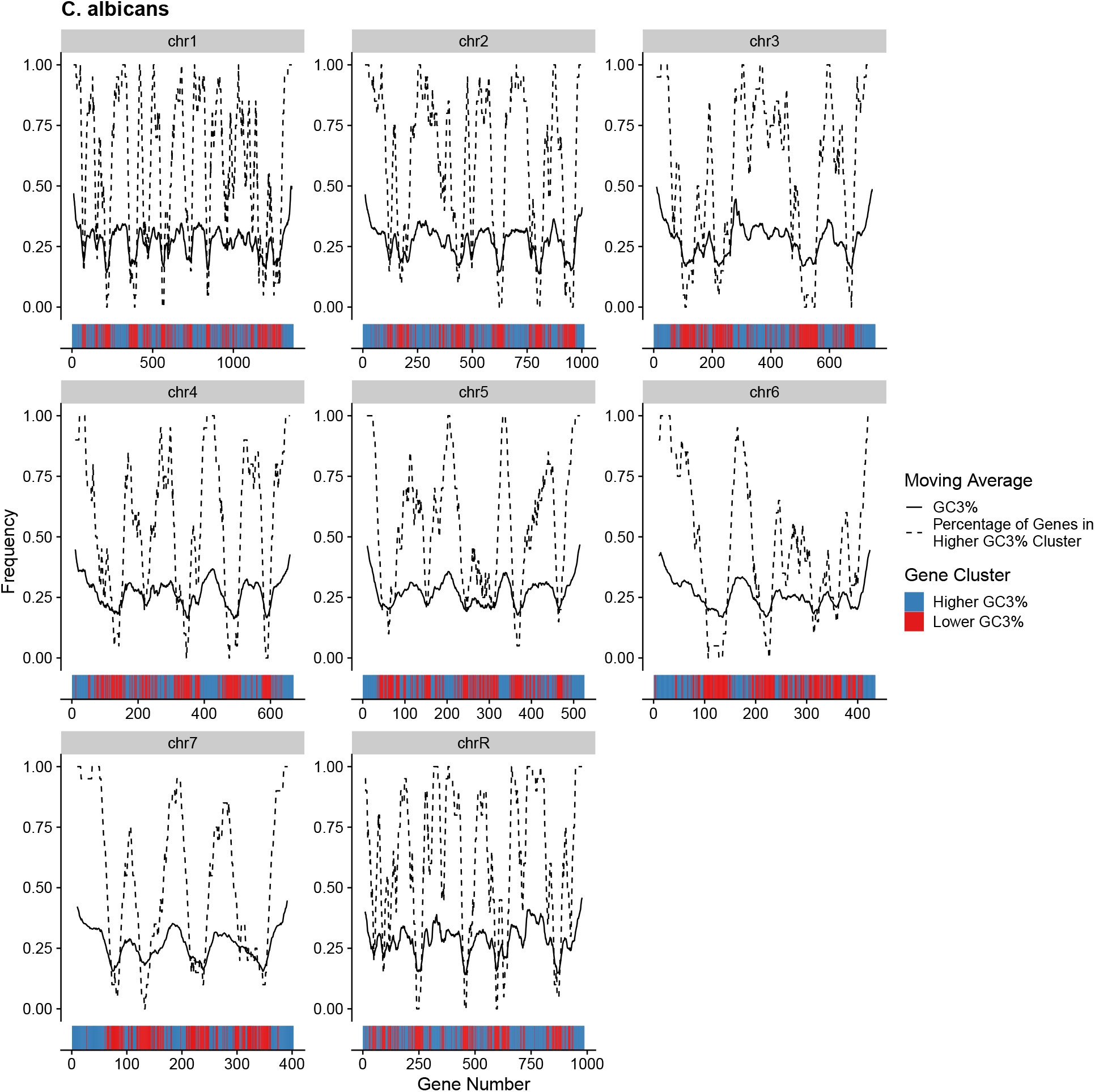
Per-gene GC3% content across all *C. albicans* chromosomes quantified as a moving average using a 20 gene sliding window (solid line). For each 20 gene window, the percentage of genes assigned to the Higher GC3% regime is also shown (dashed line). Color bars indicate the mutation regimes for Higher and Lower GC3% (blue and right, respectively).

**Figure 7:**
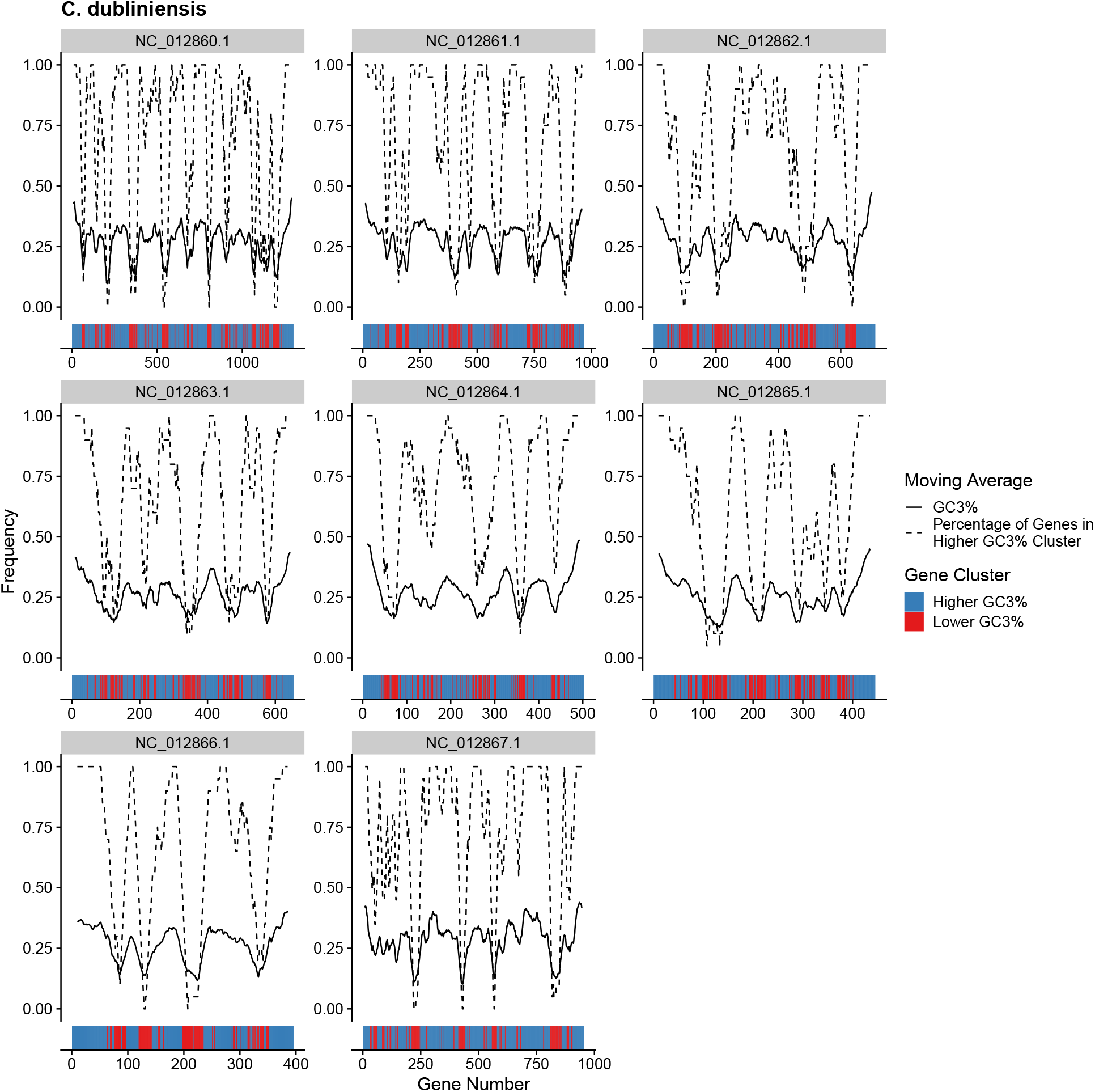
Per-gene GC3% content across all *C. dubliniensis* chromosomes quantified as a moving average using a 20 gene sliding window (solid line). For each 20 gene window, the percentage of genes assigned to the Higher GC3% regime is also shown (dashed line). Color bars indicate the mutation regimes for Higher and Lower GC3% (blue and right, respectively).

**Figure 8:**
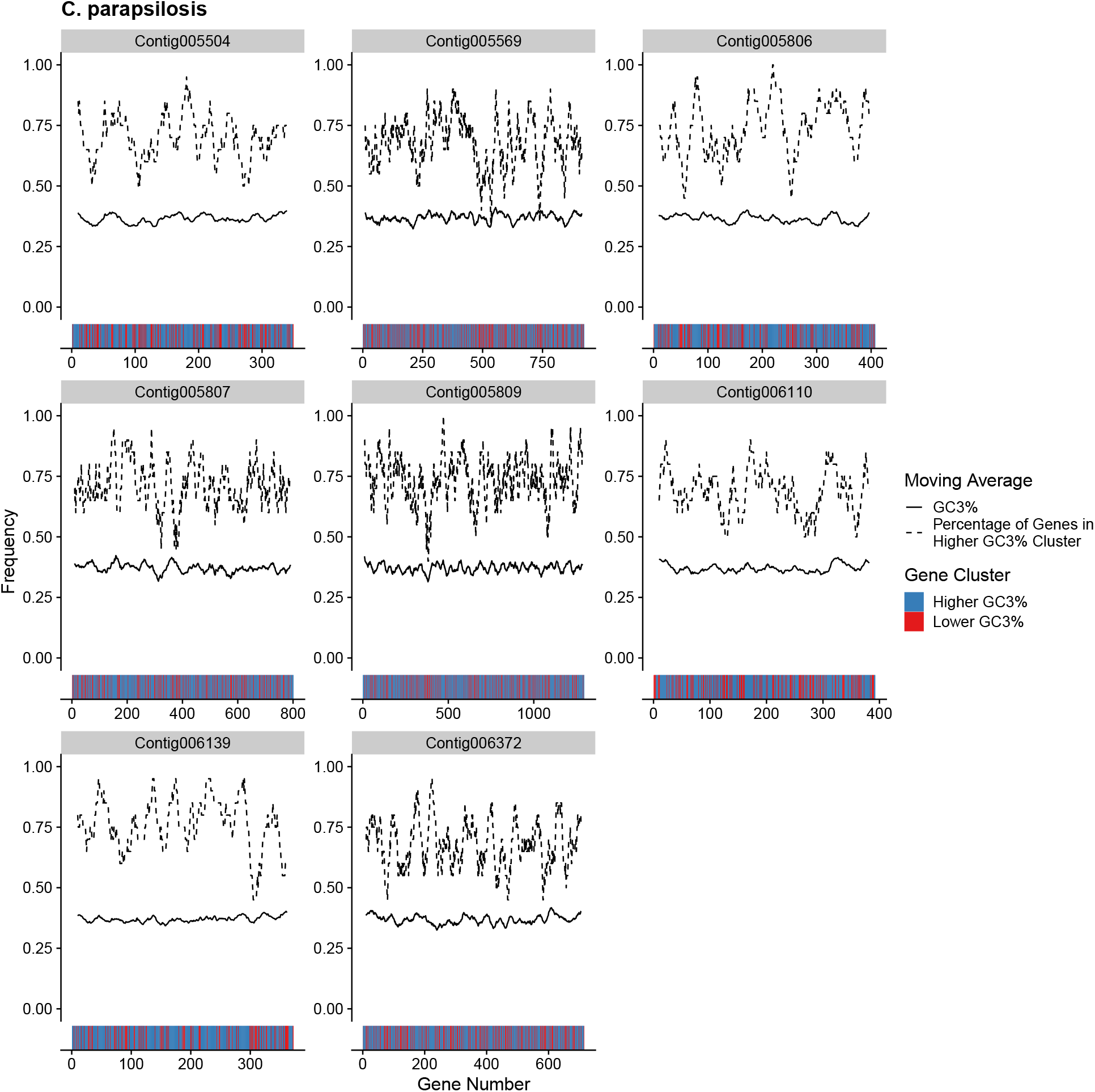
Per-gene GC3% content across all *C. parapsilosis* chromosomes quantified as a moving average using a 20 gene sliding window (solid line). For each 20 gene window, the percentage of genes assigned to the Higher GC3% regime is also shown (dashed line). Color bars indicate the mutation regimes for Higher and Lower GC3% (blue and right, respectively).

**Figure 9:**
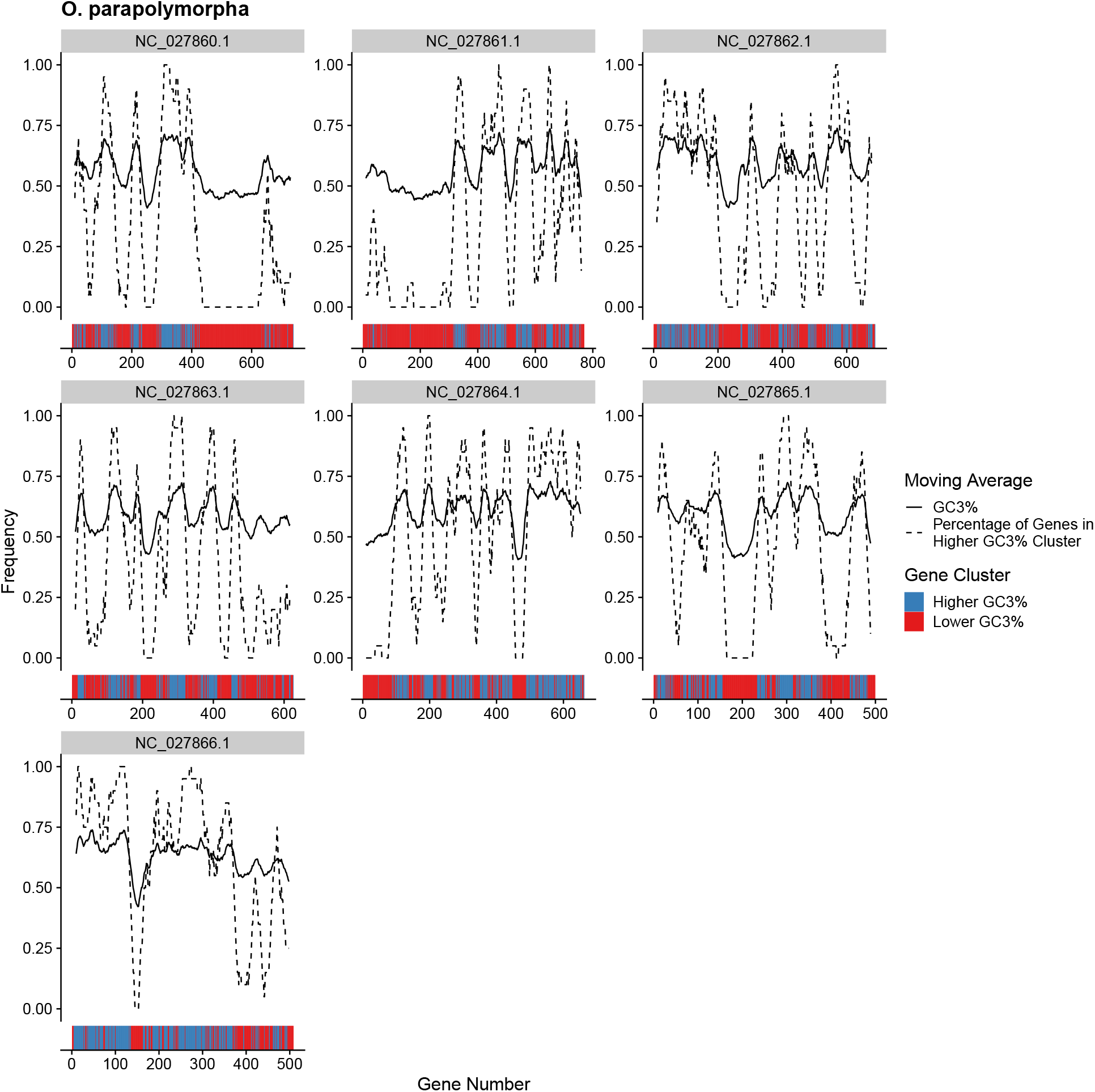
Per-gene GC3% content across all *O. parapolymorpha* chromosomes quantified as a moving average using a 20 gene sliding window (solid line). For each 20 gene window, the percentage of genes assigned to the Higher GC3% regime is also shown (dashed line). Color bars indicate the mutation regimes for Higher and Lower GC3% (blue and right, respectively).

**Figure 10:**
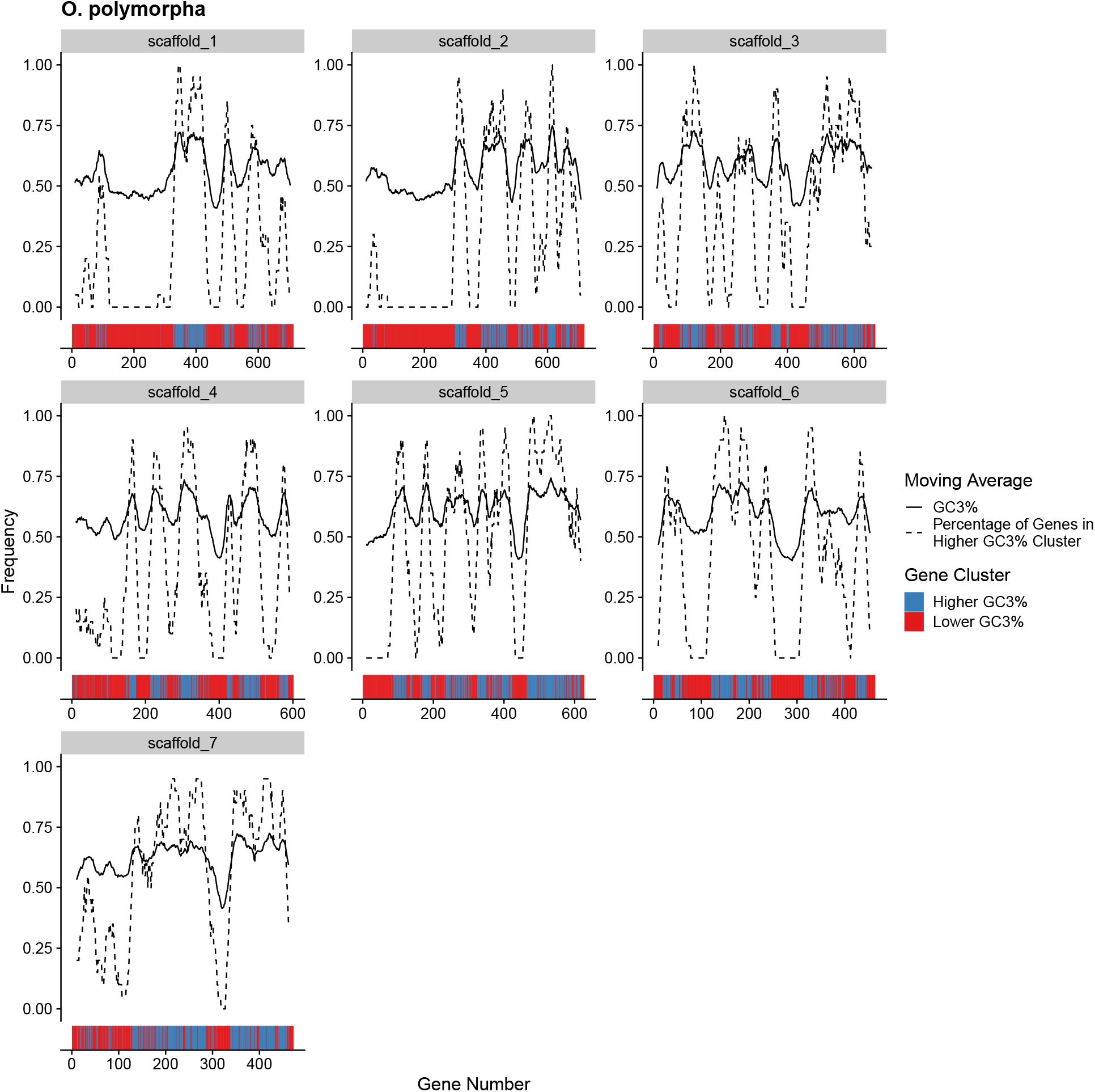
Per-gene GC3% content across all *O. polymorpha* chromosomes quantified as a moving average using a 20 gene sliding window (solid line). For each 20 gene window, the percentage of genes assigned to the Higher GC3% regime is also shown (dashed line). Color bars indicate the mutation regimes for Higher and Lower GC3% (blue and right, respectively).

**Figure 11:**
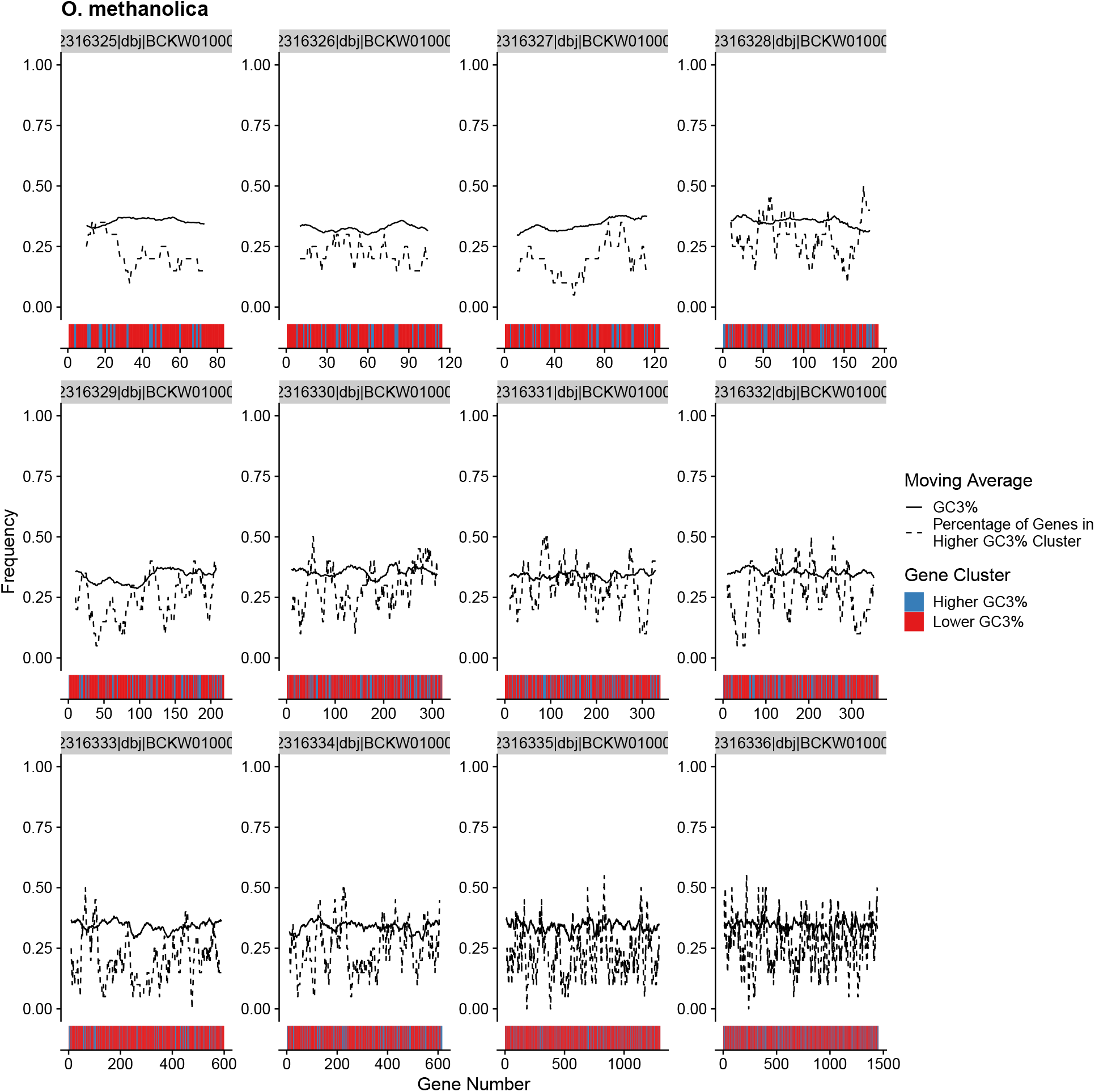
Per-gene GC3% content across all *O. methanolica* chromosomes quantified as a moving average using a 20 gene sliding window (solid line). For each 20 gene window, the percentage of genes assigned to the Higher GC3% regime is also shown (dashed line). Color bars indicate the mutation regimes for Higher and Lower GC3% (blue and right, respectively).

**Figure 12:**
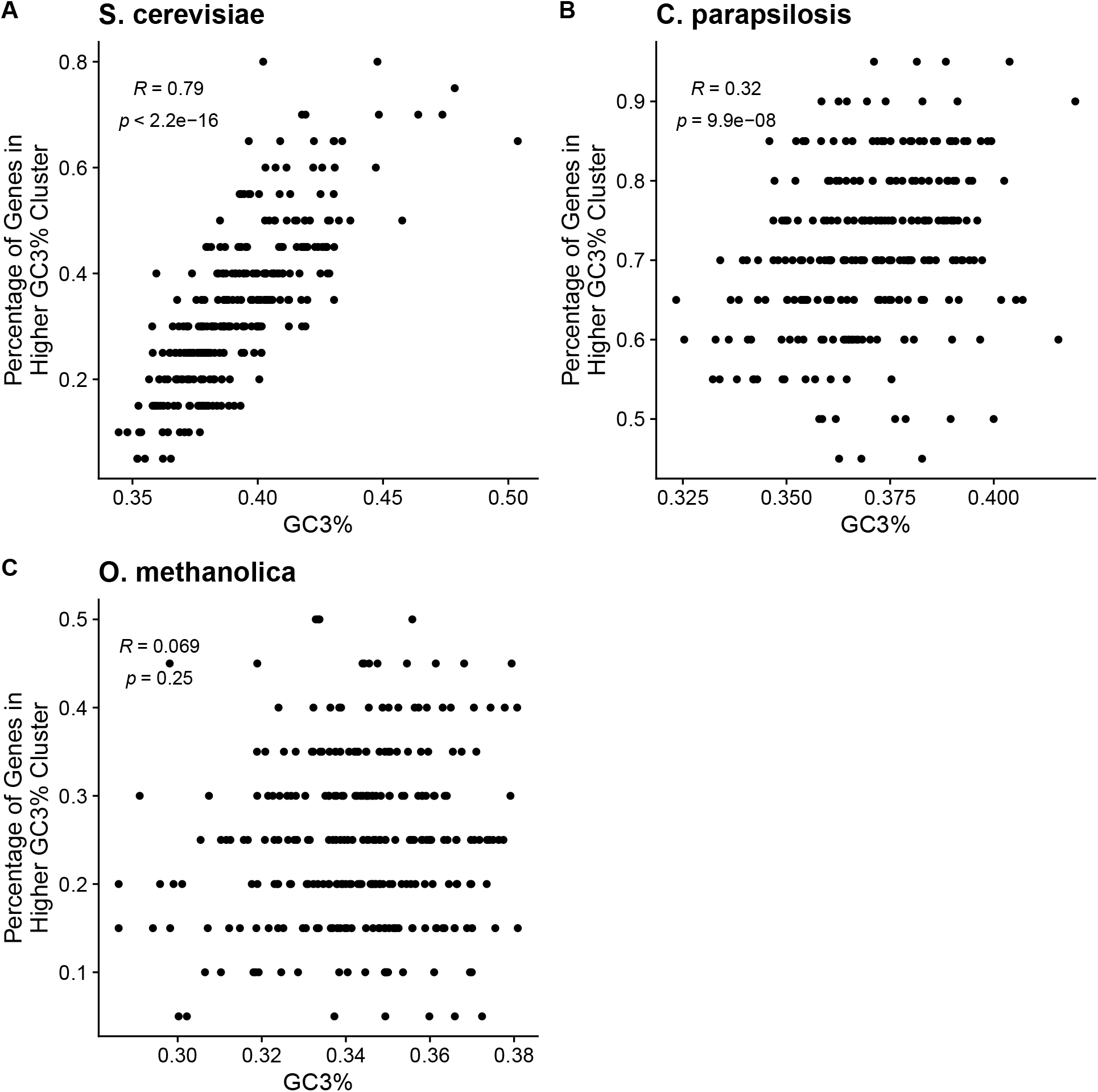
Comparing the average GC3% and percentage of genes assigned to Higher GC3% cluster using 20-gene non-overlapping windows across all chromosomes for a species. (A) *S. cerevisiae*. (B) *C. parapsilosis*. (C) *O. methanolica*.

**Figure 13:**
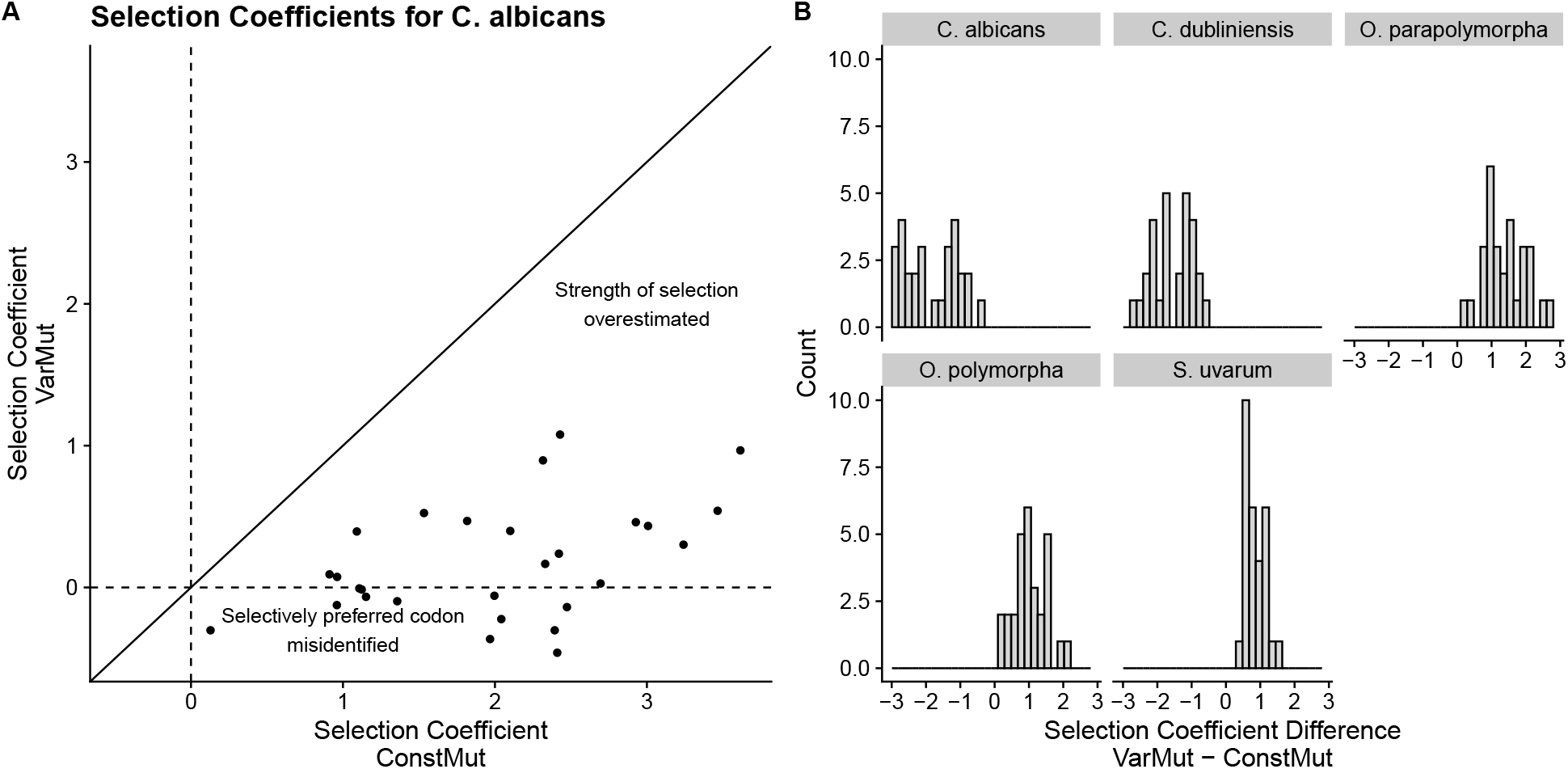
Comparison of selection coefficients estimated using the ConstMut and VarMut model. Selection coefficients are rescaled such that they represent selection of GC-ending codons relative to AT-ending codons, i.e. NNG relative to NNA or NNC relative to NNT. (A) Scatter plot showing effect of intragenomic mutation bias on misidentifying or overestimating strength of selection on GC-ending codons relative to AT-ending codons in *C. albicnas*. (B) Distribution of log fold changes of selection coefficients between VarMut and ConstMut models.

**Figure 14:**
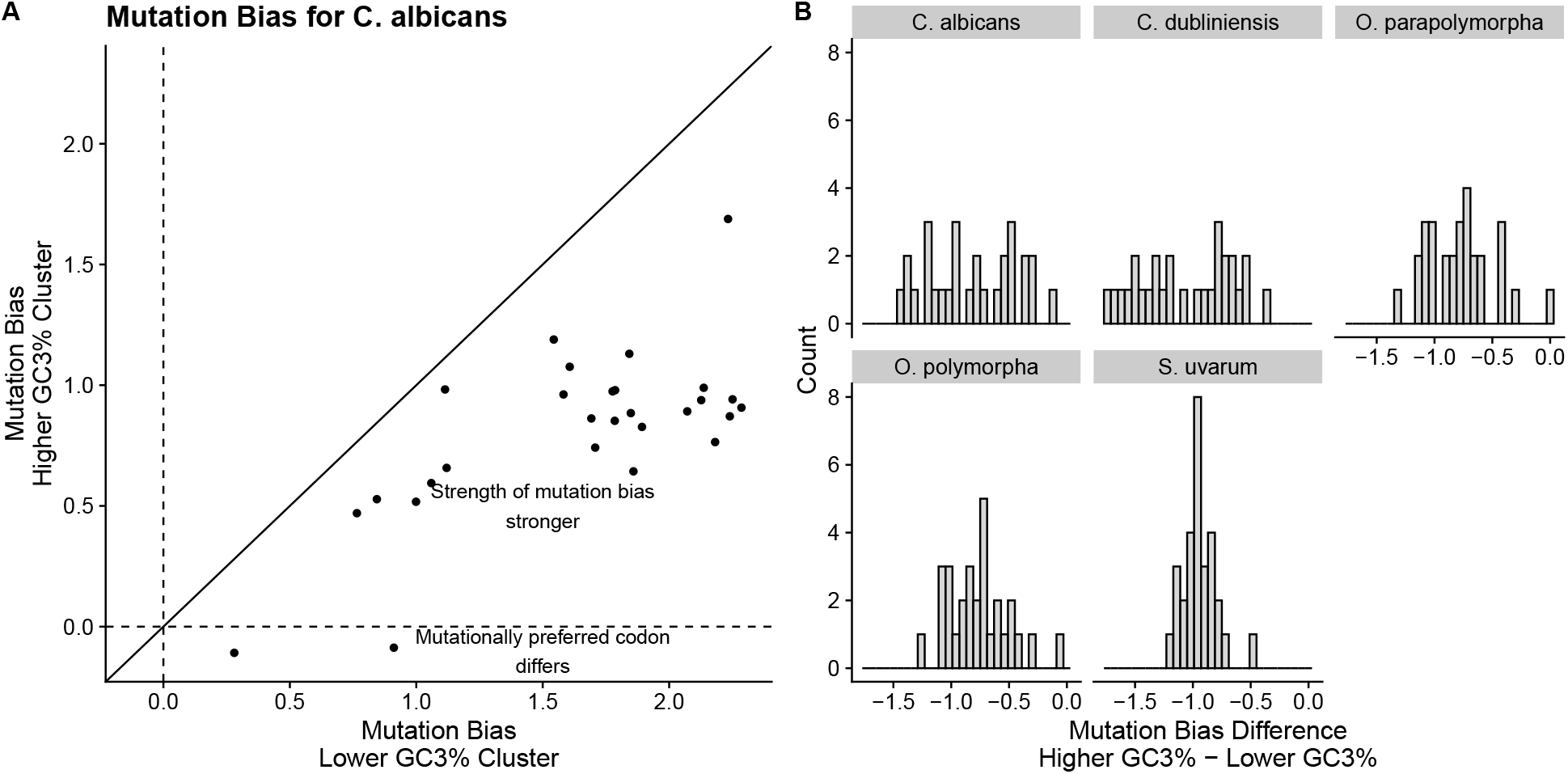
Comparison of mutation biases between the Lower and Higher GC3% clusters estimated using the VarMut model. Mutation biases are rescaled such that they represent mutation bias of GC-ending codons relative to AT-ending codons, i.e. NNG relative to NNA or NNC relative to NNT. (A) Scatter plot showing difference in mutation bias between the two clusters used in the VarMut model for *C. albicans*. (B) Distribution of log fold changes of mutation bias between clusters used in VarMut Model.

**Figure 15:**
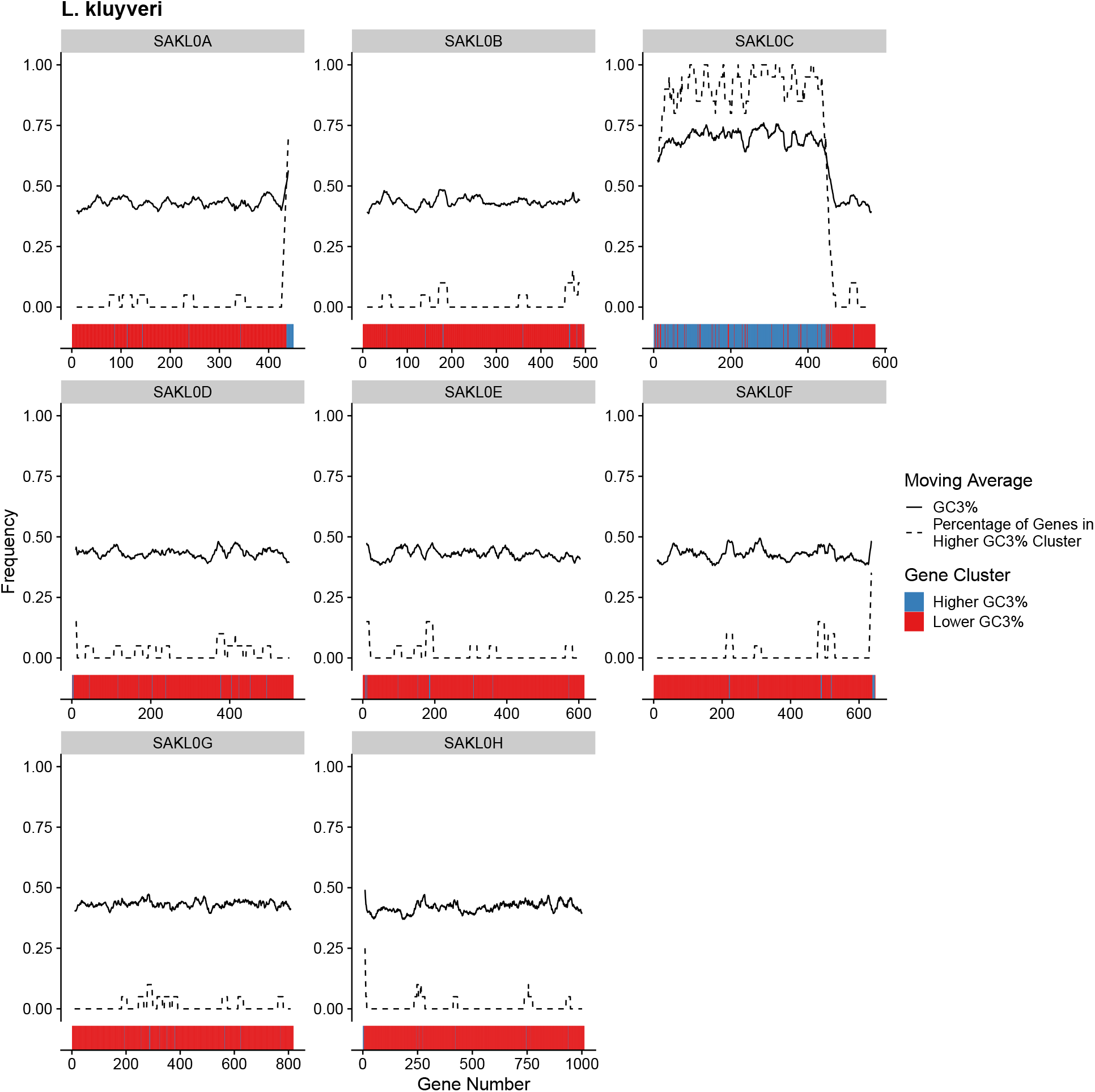
Per-gene GC3% content across all *L*.*kluyveri* chromosomes quantified as a moving average using a 20 gene sliding window (solid line). For each 20 gene window, the percentage of genes assigned to the Higher GC3% regime is also shown (dashed line). Color bars indicate the mutation regimes for Higher and Lower GC3% (blue and right, respectively). The region of high GC3% content on chromosome SAKL0C is the result of an introgression.

**Figure 16:**
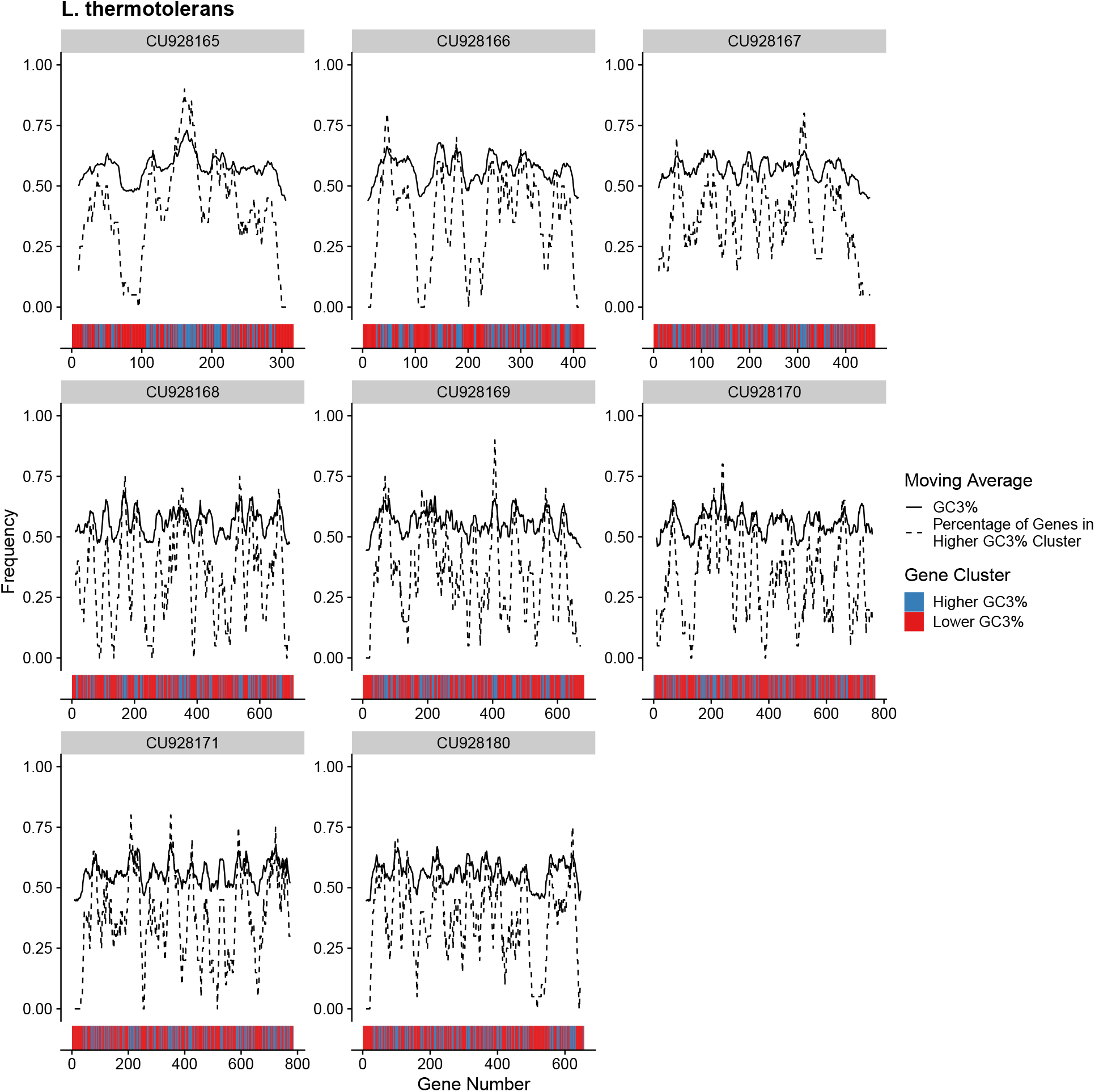
Per-gene GC3% content across all *L*.*thermotolerans* chromosomes quantified as a moving average using a 20 gene sliding window (solid line). For each 20 gene window, the percentage of genes assigned to the Higher GC3% regime is also shown (dashed line). Color bars indicate the mutation regimes for Higher and Lower GC3% (blue and right, respectively).

